# Polyketide synthase 12 is an *in vivo* essential source of novel mycolyl lipids in *Mycobacterium tuberculosis*

**DOI:** 10.64898/2026.09.25.754250

**Authors:** Gregory H. Babunovic, Sanne van der Niet, Marco T.P. Gontijo, Katherine Abrahams, Rongfeng Sun, Kiranmai Bhatt, Ella Meirav, Catherine Vilchèze, David C. Young, Lara Rosbach, Ignė Kanopaitė, Oyinda O. Adefisayo, Kirsten M. Smith, Ana María Xet-Mull, V. Grace Dellinger, Emilie Layre, Clare M. Smith, William R. Jacobs, Apoorva Bhatt, Gurdyal S. Besra, David M. Tobin, Qingyun Liu, Nicole van der Wel, D. Branch Moody

**Affiliations:** Division of Rheumatology, Inflammation, and Immunity, Brigham and Women’s Hospital, Boston, MA; Electron Microscopy Center, Amsterdam University Medical Center, Amsterdam, NL; Department of Molecular Genetics and Microbiology, Duke University School of Medicine, Durham, NC; School of Biosciences and Institute of Microbiology and Infection, University of Birmingham, Edgbaston, UK; Department of Genetics, University of North Carolina, Chapel Hill, NC; Walsall Manor Hospital NHS Trust, UK; Department of Microbiology & Immunology, Albert Einstein College of Medicine, New York, NY; Department of Genetics, Albert Einstein College of Medicine, New York, NY; University Program in Genetics and Genomics, Duke University School of Medicine, Durham, NC; Department of Integrative Immunobiology, Duke University School of Medicine, Durham, NC; Department of Microbiology and Immunology, University of North Carolina, Chapel Hill, NC

## Abstract

*Mycobacterium tuberculosis* (Mtb) is a major pathogen worldwide that infects and transmits only among humans, yet nearly all *in vivo* virulence research occurs in non-human hosts. To overcome this central challenge in tuberculosis research, we leveraged a dataset of more than 50,000 sequenced isolates to identify Mtb genes that that are functionally preserved during natural infection, disease causation, and transmission between humans. This whole-genome ranking identified polyketide synthase 12 (*pks12*) as an exceptionally *in vivo* essential gene in human tuberculosis, which we validated in zebrafish and mouse models. Though Pks12 produces a mycoketide lipid in only trace amounts, *pks12* deletion severely altered the host-facing surface and arabinoglycan architecture of Mtb. This amplified effect was explained through the discovery of mannosyl-β-1-phosphomycoketide monomycolate (MPMMM), which is synthesized from the known Pks12 product at higher abundance by antigen 85 mycolyltransferases. Thus, we used a new host-facing genomic-metabolomic-phenotypic approach to discover the functions of an *in vivo* essential Mtb gene, which controls the physical structure of the Mtb-host interface.

## MAIN TEXT

*Mycobacterium tuberculosis* (Mtb), the leading cause of death due to a single pathogen worldwide, naturally transmits, replicates, and causes disease only within human hosts (*1*, *2*). Yet unlike other infectious diseases of global significance, such as malaria and typhoid fever, no model for intentional human Mtb infection exists (*3–5*). Further, the years-long and varying tuberculosis disease course renders invasive methods needed for *in vivo* monitoring of human patients generally infeasible. Alternative research approaches have emphasized animal and cellular models to identify important genes and molecules for Mtb infection and tuberculosis disease (*6–11*). However, even keystone discoveries in this space are limited by these non-natural hosts, leaving open questions of disease relevance and limiting drug and vaccine target investigation. After the failure of several recent tuberculosis vaccine trials (*12–15*), devising better approaches to identify *in vivo* essential targets in human tuberculosis is one of the greatest unmet needs for reducing disease burden.

### *pks12* is *in vivo* essential in humans

We previously measured positive selection to identify genes that are liabilities or otherwise favored for change in Mtb, especially in the face of historically recent antibiotic pressures (*16*, *17*). To generate a new measurement of *in vivo* gene essentiality in humans, we took the opposite approach of measuring purifying selection—searching for genes that are strongly preserved during the Mtb infectious lifestyle in human populations, and loss of which could therefore be inferred to cause an evolutionary dead end. We reasoned that since the particular Mtb organisms in patient sputum isolates have successfully transmitted to, infected, and caused lung disease within a human host under natural epidemic conditions, they must have functionally preserved the genes required to do so. The recent expansion of Mtb clinical genomic datasets to a massive scale allowed us to measure this preservation through multiple metrics of purifying selection.

First, we identified genes across 51,229 sequenced clinical isolates that did not undergo truncation via mutational formation of premature stop codons. Genes that are essential *in vitro* during culture typically sustain few such truncations; this is expected, as mutations in these genes can disable fundamental Mtb cellular functions independently of any specific interaction with the host (Fig. 1A). Other genes have important host-oriented roles during infection beyond basic mycobacterial cell survival and division, including genes whose deletion does not affect growth in media but crucially and counterintuitively impairs survival in humans. Indeed, we found that 10% of *in vitro* nonessential genes also sustained zero truncations *in vivo* (Fig. 1B, Table S1). While for shorter genes this could be due to random chance, the probability of truncating mutations increases with gene length. Thus, we used this statistical metric of purifying selection to rank-order genes, with the longest being the top *in vitro* nonessential yet *in vivo* essential candidates (Fig. 1B, bottom right, Fig. S1).

**Fig. 1.**
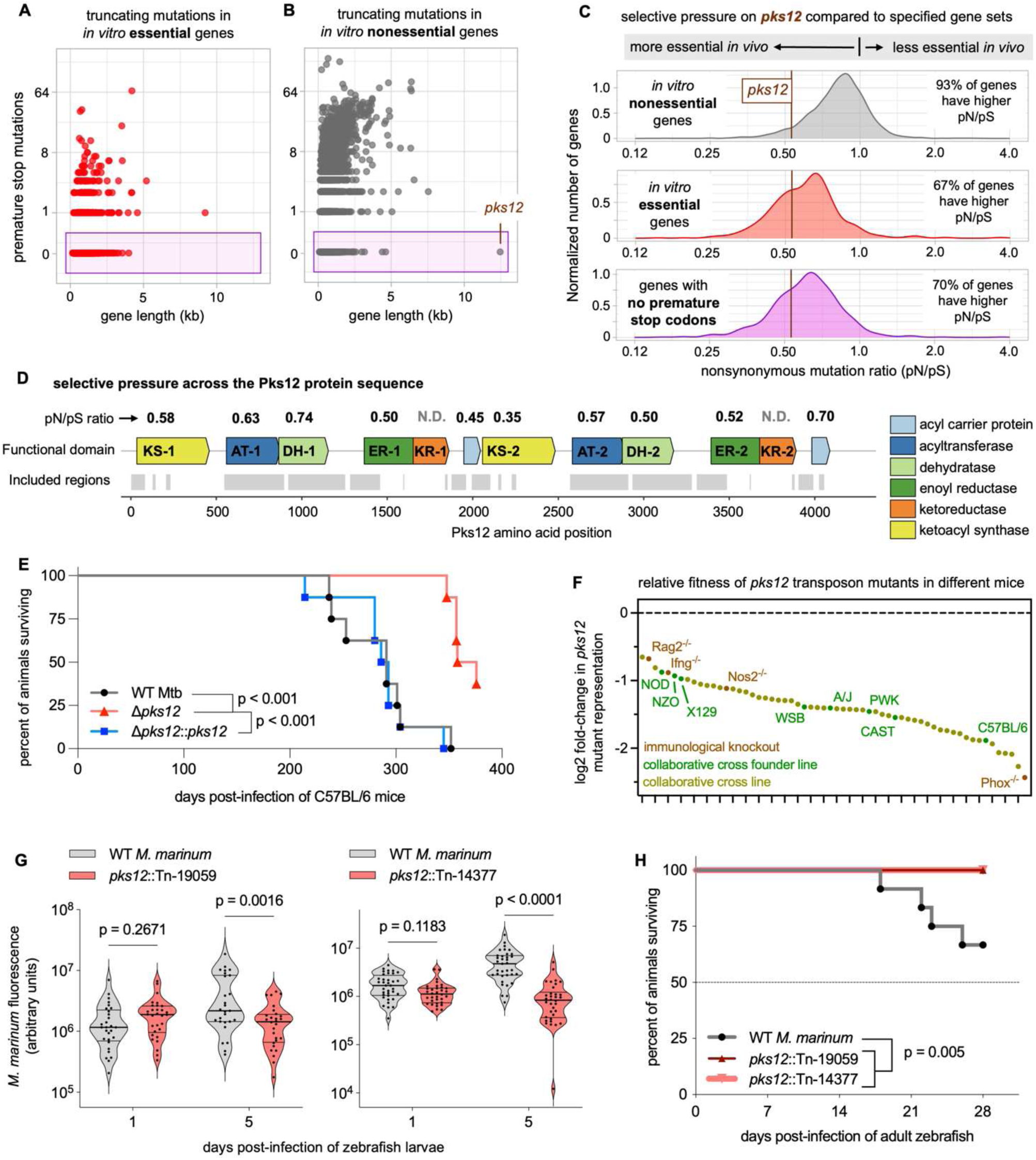
***pks12* is indispensable in humans and important in animal models.** (**A-D**) Analysis of genomes from 51,229 sequenced Mtb isolates. (**A,B**) Number of premature stop codon mutations per measured *in vitro* essential (**A**) or nonessential (**B**) gene, based on published essentiality calls. (**C,D**), Level of purifying (pN/pS < 1) versus positive (pN/pS > 1) selection for different sets of genes, with the pN/pS of *pks12* represented by vertical lines (**C**), and for each measurable domain of the Pks12 protein (**D**); regions not measured in (**D**) were too similar to another region to accurately call variants. (**E**) Survival of C57BL/6 mice following aerosol infection with approx. 100 colony-forming units of Mtb. (**F**) Fitness of Mtb transposon mutants in *pks12* during pooled whole-genome library infections across mouse genetic backgrounds, calculated from published data. (**G**) Bacterial burden, as quantified by mCerulean fluorescence, in zebrafish larvae infected with either WT *M. marinum* or two independent *pks12* transposon mutants; each point represents one larva. (**H**) Survival of adult zebrafish infected with indicated *M. marinum* strains. Statistics were calculated with log-rank tests (E and H, with Benjamini-Hochberg corrected multiple testing in E), or with a mixed model with Šídák’s multiple comparisons test (G).

These genes are of particular interest because they can be specifically required for host-dependent Mtb processes in humans, such as stress response or transmission. We found that the longest zero-truncation and therefore most evidently *in vivo* essential Mtb gene was the *in vitro* nonessential gene encoding polyketide synthase 12 (*pks12*, Fig. 1B, Fig. S1). Validating this new *in vivo* essential gene discovery approach, *pks12* was also under exceptionally strong purifying selection by the independent and conventional metric of nonsynonymous variant ratio (pN/pS), which is inversely related to a gene’s *in vivo* essentiality. The *pks12* ratio was lower than that of the overwhelming majority of other nonessential genes, and a large majority of *in vitro* essential genes, as well as all other genes with no measured premature truncations (Fig. 1C). This signal of *in vivo* essentiality (pN/pS well below 1) was present across every measurable domain of the Pks12 protein (Fig. 1D), supporting the essentiality of Pks12 enzymatic function—which requires all domains (*18*). Thus, despite its lack of effect on growth in standard media (*19*), *pks12* is *in vivo* essential in humans by multiple metrics.

### *pks12* **deletion attenuates mycobacteria in** *in vivo* **models**

Next we validated this human-focused gene discovery approach using animal models, with disease severity and bacterial growth as proxies for fitness in humans. Compared with a matched wild type (WT) Mtb strain, deletion of *pks12* slowed the lethality of aerosol infection with 100 colony-forming units (cfu) of Mtb in C57BL/6 mice. This effect was highly significant and fully reversed upon genetic complementation, while appearing at a relatively late 200-day timepoint following durable Mtb-host interaction (Fig. 1E). Standard mouse lines are inbred and can produce phenotypes that may be unique to a single host strain; in contrast, bacterial isolates in our evolutionary analysis infected globally diverse human populations. To test the *in vivo* effects of *pks12* across genetically diverse hosts, we leveraged a published mixed infection experiment to examine *pks12* effects in 60 genetically defined and distinct mouse lines, including collaborative cross mice, collaborative cross founders, and immune deletions (*6*). In a pooled genome-scale transposon mutant library, *pks12* is one of only 10 genes among 4,111 tested for which insertion mutants are significantly less fit (Q < 0.05) across every measured mouse genetic background (Fig. 1F, Table S2), demonstrating high penetrance across hosts. *Phox*^-/-^ mice showed strongest attenuation while *Rag2*^-/-^, *Ifng*^-/-^, and NOD showed the least attenuation, pointing to potential interactions with immunity.

Mice are a valuable and tractable model for Mtb infection, but often fail to recapitulate specific certain aspects of tuberculosis disease, such as the formation of distinct granulomas (*20*). To examine the role of *pks12* during mycobacterial infection in a granuloma-forming model, and to further diversify evidence on *pks12* importance, we turned to *Mycobacterium marinum* infection of zebrafish. We identified and confirmed two independent *M. marinum* transposon mutants in *pks12*, both of which disrupted Pks12 enzymatic function (Fig. S2A-C). We infected zebrafish larvae with fluorescently labeled versions of these transposon mutants and observed reduced bacterial burden following five days of infection compared to the WT strain (Fig. 1G, Fig. S2D). Zebrafish larvae rely on innate immunity; as in mammals, adaptive immunity develops later. To examine the role of *pks12* in fully immunocompetent animals and during longer-term infections where more mature granulomas are developed, we infected adult zebrafish. The *pks12* transposon mutants failed to kill adult zebrafish after 4 weeks of infection, while WT *M. marinum* significantly decreased host survival (Fig. 1H). Whereas it was previously unknown if *pks12* controlled mycobacterial growth or host survival during infection, loss of this top ranked human *in vivo* virulence gene attenuated Mtb across all tested infection models.

### *pks12* deletion interferes with cell envelope and capsular homeostasis

Pks12 is a 12-domain enzyme that synthesizes mycoketide (Fig. 1D), the methyl-branched acyl chain of the atypical phospholipid mannosyl-β-1-phosphomycoketide (MPM) (*18*, *21*). MPM is a trace molecule of unknown function in Mtb, present at approximately 1 part per million of total lipid mass (*21*). Because of the strong survival effects across all tested models, we hypothesized that this trace metabolite impacts mycobacterial survival *in vivo* through processes that are amplified by specific receptor recognition, including immune mechanisms, rather than broader and direct changes in cell envelope structure.

To test the hypothesis of minimal cell envelope changes, we performed electron microscopy (EM) on Mtb stained with osmium tetroxide and potassium ferrocyanide to preserve mycobacterial envelope surface morphology and visualize the native glycan capsular layer (*22*) (Fig. S3). Opposite to expectations, scanning EM (SEM) revealed that the Δ*pks12* mutant exhibited common and pronounced longitudinal cell surface indentations, compared to WT and complemented strains (Fig. 2A-C, cyan arrows). These striking ’canyon-like’ indentations extended through all layers of the cell envelope, including inner plasma membrane, outer mycomembrane, and capsule, as shown by transmission EM (TEM) analysis of cryosectioned bacteria (Fig. S4). We also observed smaller mushroom-like exophytic punctate structures on the surface of the Δ*pks12* mutant by SEM (Fig. 2D, magenta arrows), and by TEM identified similar electron-dense structures extending from the capsule in a significantly higher proportion of Δ*pks12* cells than in matched WT or complemented strains (Fig. 2E-F, purple arrows).

**Fig. 2.**
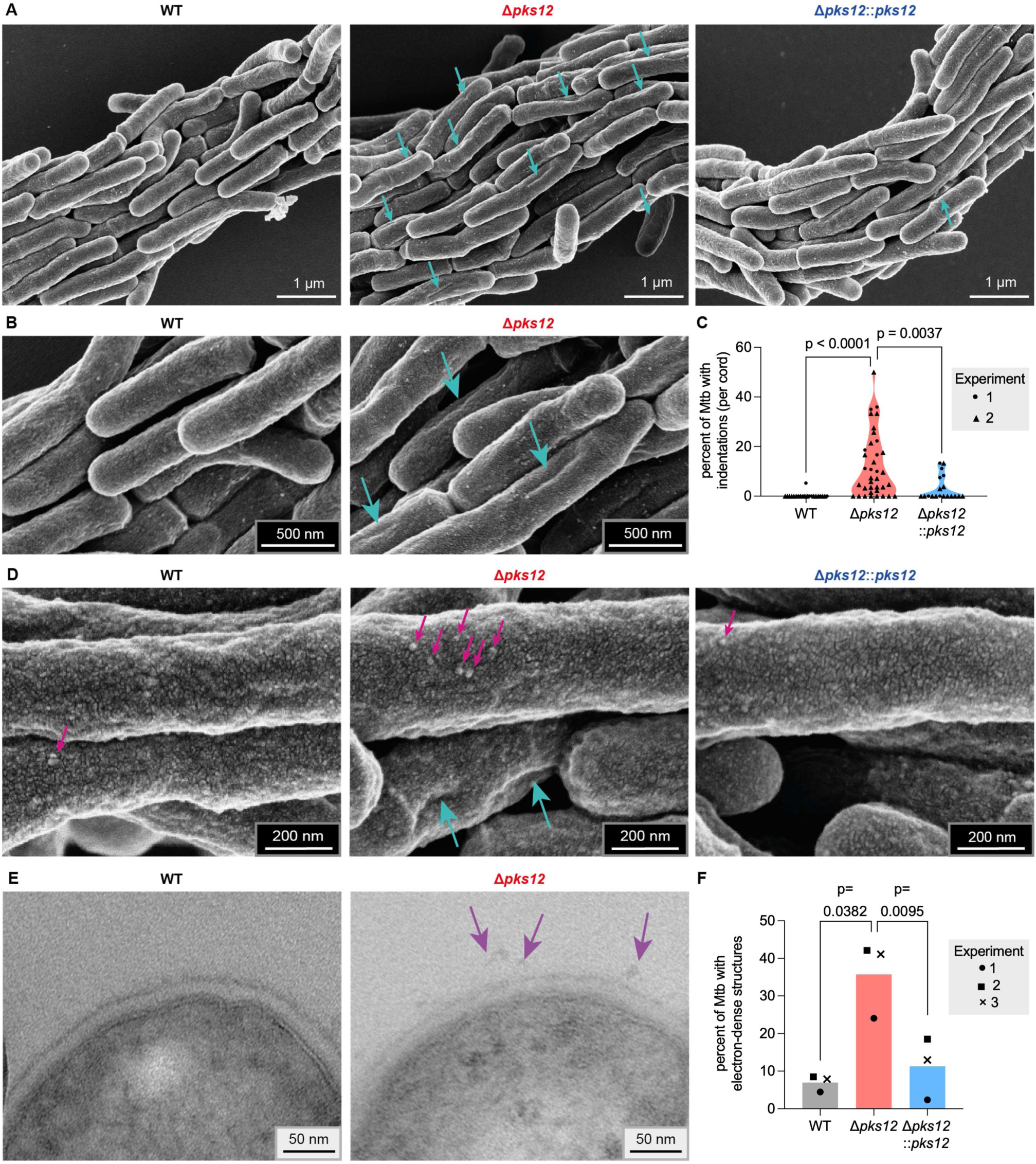
Deletion of *pks12* disrupts Mtb cell envelope morphology. (**A**) Representative SEM images of WT, Δ*pks12,* and complemented Mtb in native cord structures; cyan arrows indicate longitudinal indentations. (**B**) Magnified selected images from (A). (**C**) Quantification of indentations on the Mtb surface, from two independent experiments; statistics were performed using the Kruskal-Wallis method with Dunn’s multiple comparisons test. (**D**) Representative SEM images of the surface of WT, Δ*pks12* and complemented Mtb; magenta arrows indicate puncta on the surface and cyan arrows indicate indentations. (**E**) Representative TEM images of WT and Δ*pks12* Mb; purple arrows indicate electron-dense structures protruding from the capsular layer. (F) Quantification of electron-dense structures attached to the capsular layer of WT, Δ*pks12,* and complemented Mtb, from three independent experiments; statistics were performed using a matched one-way ANOVA with Dunnett’s multiple comparisons test.

### *pks12* **deletion mislocalizes cell envelope glycans**

Pks12 is a cytosolic enzyme producing a trace phospholipid, and mycobacterial phospholipids are typically found in the inner membrane (*23*). We were therefore surprised that deleting *pks12* had effects throughout the diderm cell envelope, clearly extending to the cell surface. This outcome could be due to downstream effects of MPM on more abundant molecules found in multiple envelope compartments. Since MPM chemically resembles the known periplasmic mannose donor decaprenylphosphoryl mannose (*24–26*), we hypothesized that it may serve as an alternative mannose donor for a mannolipid or glycan (Fig. 3A-B). One such molecule found across compartments is arabinomannan (AM), which is produced on the inner membrane in a lipidated form and is present in the capsule following enzymatic processing (*27*). Most of the mannose in lipoarabinomannan (LAM) and AM is sourced from lipid-linked donors (*28*); if MPM serves as a donor for an alternative mannose transfer pathway, this could cause *pks12*-dependent changes in AM and LAM.

**Fig. 3.**
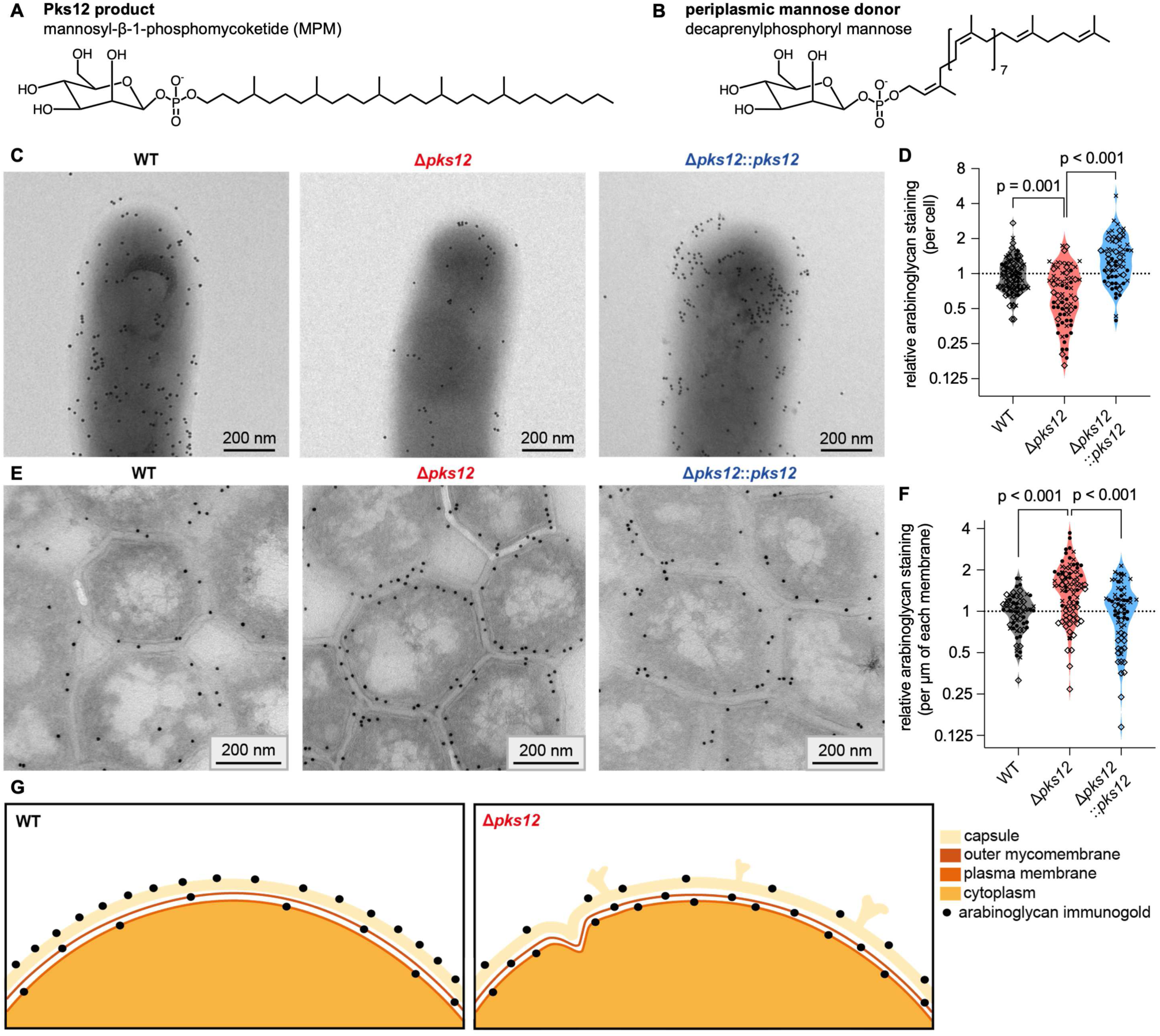
Deletion of *pks12* disrupts Mtb envelope arabinoglycans. (**A**) Structure of MPM. (**B**) Structure of the known Mtb polyprenyl mannose donor. (**C**) Representative TEM images of intact WT, Δ*pks12,* and complemented Mtb, grown in rich medium, after CS-35 immunogold staining for surface arabinoglycans. (**D**) Quantification of of CS-35 immunogold on intact Mtb as shown in (C); data points represent individual bacteria. (**E**) Representative TEM images of cryo-sections from WT, Δ*pks12,* and complemented Mtb, grown in rich medium, after CS-35 immunogold staining for arabinoglycans. (**F**) Quantification of of CS-35 immunogold on Mtb cryosections as shown in (E), per µM of membrane; data points represent individual bacteria. (**G**) Schematic representation of CS-35 immunogold distribution and morphological changes across the cell envelope of WT and Δ*pks12* Mtb. Data in (D) and (F) are from three independent experiments, indicated by different symbols; statistics were performed using linear mixed-effects models with Tukey’s multiple comparisons test.

To measure changes in AM across the cell envelope we performed immuno-gold labeling of arabinan-containing glycans—including both LAM and AM—using the CS-35 monoclonal antibody that specifically recognizes a hexa-arabinosyl motif found on these molecules (*29*). Staining of intact cell surfaces was significantly reduced in the Δ*pks12* mutant relative to WT and complemented strains, specifically on Mtb grown in a rich medium (Fig. 3C-D, Fig. S5A). To determine whether this was due to reduced production of lipoarabinomannan (LAM) we stained cryosections, finding that the Δ*pks12* mutant accumulated significantly more arabinoglycans on the inner membrane (Fig. 3E-F). This suggested that *pks12* deletion, rather than reducing AM or LAM synthesis, instead reduced AM present on the cell surface while increasing LAM at the inner membrane. Further investigation of cords (aligned clusters) (*30*) of Mtb cells revealed that Δ*pks12* bacteria were significantly more likely to accumulate arabinoglycans at the outermost layer of the cord adjacent to the culture medium, while WT and complemented Mtb accumulated more of these glycans at the interface in between bacteria (Fig. S5B-D). Overall, reduced cell surface arabinoglycan and uneven capsular structures suggest disrupted capsular integrity; combined with deep envelope indentations and changes in intracellular glycan distribution, these findings constitute a multifaceted impact of deleting *pks12* and disrupting MPM biosynthesis (Fig. 3G). This outcome contradicted our initial hypothesis, and suggested that these unexpected changes in the encapsulated Mtb cell surface—which forms the host-pathogen interface *in vivo*—are key to *pks12* function and understanding why it is essential during human infection.

### Most MPM is mycolated to form a novel lipid

The biochemical role of MPM, and therefore mechanistically how *pks12* disruption impacts the cell envelope and causes a substantial evolutionary constraint in human hosts, remained unknown. Though the polyprenyl-like structure of MPM (Fig. 3A,B) and our functional studies hint at a role in mannose transfer, this is not proven by downstream effects on AM and LAM alone. We therefore turned to untargeted metabolomic analysis, which can uncover functional mechanistic consequences downstream of genetic disruptions. As mycobacteria are unusually lipid-rich, lipid-focused metabolomics (lipidomics) has been an especially useful tool for Mtb (*31–37*). We performed unbiased lipidomics in WT, Δ*pks12*, and complemented strains of Mtb using high performance lipid chromatography-electrospray ionization mass spectrometry (HPLC-ESI-MS). As expected, MPM was absent specifically in the Δ*pks12* mutant (Fig. 4A). We hypothesized that, given the severity of cell envelope phenotypes in EM experiments and the low abundance of MPM itself, MPM may be essential for the biosynthesis of other more abundant lipids in Mtb. Untargeted analysis of lipid ions identified by automated peak-picking software failed to identify any molecules apart from MPM that were entirely lost after *pks12* deletion (Fig. S6, Table S3).

**Fig. 4.**
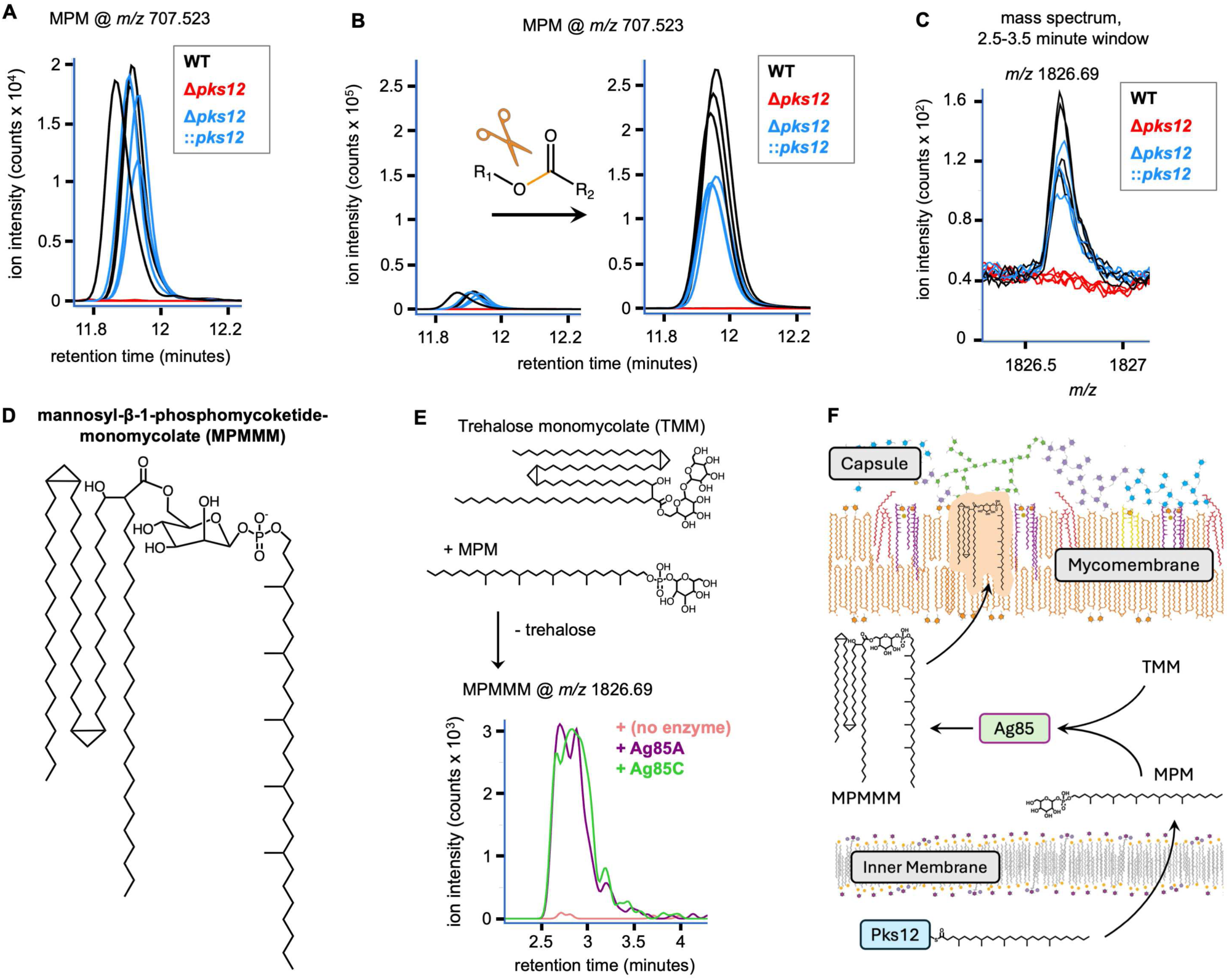
The primary product of Pks12 is mycolated MPM. (**A**) Reversed phase HPLC-ESI-MS chromatograms for MPM across strains with differing *pks12* status. (**B**) Reversed phase HPLC-ESI-MS chromatograms for MPM before (left) and after (right) saponification of total Mtb lipids. (**C**) Mass spectrum from an early window of normal phase HPLC-ESI-MS across strains. (**D**) Structure of the most abundant form of MPMMM; linkage at the C6 position of mannose is presumed based on mycolyltransferase preference. (**E**) Normal phase HPLC-ESI-MS chromatograms for MPMMM following incubation of substrates with Ag85 enzymes. (**F**) A model for MPMMM production and presumed localization in Mtb.

Some large lipids are poorly ionizable and thus challenging to detect by MS; to account for these, we chemically released more easily detected fatty acids through saponification and performed HPLC-ESI-MS analysis. Surprisingly, this revealed that a large pool of MPM—far more abundant than the free pool in Mtb—is esterified to form one or more larger molecules (Fig 4B). Whereas the MPM phospholipid is polar, fractionation showed that this MPM-ester-linked material is hydrophobic, segregating into nonpolar solvents and eluting very early during normal phase HPLC (Fig. S7). Review of mass spectra in this early range identified an alkane family of *pks12*-dependent lipids, with the most intense ions having a nominal monoisotopic mass/charge ratio (*m/z*) of 1826 (Fig. 4C, Fig. S8A). These ions did not match any *m/z* values in the MycoMass database, suggesting that their shared lipid structure was unknown (*32*, *36*). They were also not identified in our previous analysis, likely due to high mass and poor ionization precluding their identification by automated peak-picking software. Collision-induced dissociation mass spectrometry (CID-MS) of an abundant representative ion produced fragments matching mycolic acid (*m/z* 1136.16) and deoxy-MPM (*m/z* 689.513), along with diagnostic fragments for a mycolate α-chain (*m/z* 395.390), phosphomycoketide (*m/z* 545.469), and a through-ring cleavage product characteristic of the β-1 linkage in MPM (*m/z* 587.479) (Fig. S8B) (*38*, *39*). These data identified this *pks12*-dependent lipid class as a previously undiscovered molecule, MPM monomycolate (MPMMM) (Fig. 4D). Though the soil-dwelling saprophyte *Mycobacterium smegmatis*—which does not encode *pks12*—produces a different mycolated phospholipid (*40*), to our knowledge no such molecule has been described in Mtb or any other pathogen.

### MPMMM is synthesized by Antigen 85 mycolyltransferases

The mycolated nature of MPMMM suggested a role in the mycolate outer membrane (mycomembrane) of Mtb. As *pks12* and MPM are nonessential *in vitro* and not found in all mycobacteria, MPMMM cannot be an essential precursor or carrier for broadly shared mycomembrane components. We therefore hypothesized that MPMMM is instead the product of an alternate biosynthetic pathway using the same mycolyl donors as known mycomembrane components. *pks12*—along with an unknown phosphotransferase, and the nearby mannosyltransferase gene *ppm1*—can account for MPM biosynthesis in the cytoplasm (*18*, *21*). We hypothesized that MPMMM is generated downstream of MPM by mycolyl transfer, possibly via antigen 85 (Ag85) enzymes which are known to transfer mycolic acid from trehalose monomycolate (TMM) to arabinose- and trehalose-containing substrates. Through *in vitro* enzymology, we found that Ag85A and Ag85C both efficiently catalyze mycolic acid coupling to MPM to generate MPMMM (Fig. 4E, Fig. S9).

While *pks12* is essential for MPM and MPMMM production, it is unclear *a priori* whether MPMMM is the major lipid effector downstream of Pks12. Although no method can fully rule out other MPM partners or substrates in Mtb, analysis of the Δ*pks12* lipidome failed to identify other lipids lost with *pks12* deletion (Fig. S6). Far more MPM signal is produced following saponification than is directly detectable, highlighting MPMMM as the major accumulating product of this pathway (Fig. 4B). These saponification experiments detected MPM release from one discrete retention time range that resolves highly hydrophobic molecules, rather than diffuse positions in the HPLC run, supporting the identification of MPMMM as the main or sole esterified MPM product (Fig. S7D). Combined, these observations strongly suggest that MPMMM is the most abundant product of the Pks12-nucleated pathway, and thus is the best candidate for mediating the essential *pks12* function during natural human tuberculosis and virulence effects in animal models.

The unanticipated role of Ag85 enzymes, which are secreted to the periplasm and Mtb surface (*41*), supports a new stepwise export-dependent biosynthetic pathway for MPMMM—like those seen for sulfoglycolipid and trehalose dimycolate (*42–44*) (Fig. 4F). This is consistent with the presumed localization of MPMMM to the cell surface, where its absence could cause the canyon-like clefts, exophytic clusters, and broad mislocalization of arabinoglycans observed in the Δ*pks12* mutant (Fig. 3G). Although future work is needed to define the detailed mechanisms behind glycan remodeling, the MPMMM chemical structure provides potential clues, as it is comprised of a mycolyl lipid and a phosphorylated pseudopolyisoprenol: the resulting amphipathic molecule could control the interface between the hydrophobic outer membrane and the polar, loosely adherent capsular glycans. Overall, using a discovery method geared to identify the most functionally host-dependent virulence factors of Mtb in humans, we identified a new cell envelope biosynthetic pathway that broadly controls Mtb’s glycan-rich structural boundary with the host.

Given that Mtb only infects and transmits in humans and remains a major cause of death, improved human-focused approaches for understanding virulence mechanisms are urgently needed. The key to this discovery of a human-host-dependent virulence factor was analysis of massive genomic datasets from clinical isolates, combining novel and conventional metrics to identify genes essential to Mtb during the *in vivo* gauntlet of initial infection, disease causation, aerosolization, and re-entry into the airways. Beyond *pks12*, this study also generated ranked lists of *in vitro* nonessential genes with a high likelihood of *in vivo* essentiality, including those with zero premature truncations (Table S1).

These include known virulence and stress response factors, such as *ppsB* and *relA*, as well as genes of unknown function that could represent additional virulence pathways required for survival within or between human hosts. Our findings also highlight the importance of undiscovered metabolites in one of the world’s most extensively studied pathogens: more than 100 years after the unusually lipid-rich nature of Mtb was first noted, approximately half of its detectable lipidome remains unannotated (*31*, *37*). The combined genomic-metabolomic-phenotypic approach used in this work can be rapidly applied to prioritize genes for study and understand which gene products and metabolites are most important in the human host, directly informing the design of vaccines and therapeutics targeting factors Mtb cannot afford to lose.

## ACKNOWLEDGMENTS

We thank Christine Cosma and Lalita Ramakrishnan for their generosity in providing a sequenced *M. marinum* transposon library. We also thank Tan-Yun Cheng, Yashodhan Nair, Kyu Rhee, and Adriaan Minnaard for insightful comments throughout the experimental preparation of this work, and Adriaan Minnaard for providing synthetic MPM. This work was funded by the following NIH grants: U19AI162584, P01AI201075, R01AI165573, and K99AI96044.

## METHODS

### Ethics statement

All animal husbandry and experiments were performed in accordance with National Institutes of Health standards for the care and use of animals. Zebrafish (*Danio rerio* *AB) protocols were approved by the Duke University Animal Care and Use Committee under protocol A049-23-03 and #9. Mouse protocols were approved by the Albert Einstein College of Medicine Institutional Animal Care and Use Committee.

### Genomic analysis of purifying selection

A global collection of 51,229 publicly available *Mycobacterium tuberculosis* complex (MTBC) genomes was previously curated (*16*). Briefly, raw sequencing reads were quality trimmed using Sickle (*45*), and reads longer than 30 bp with Phred quality scores >20 were retained for subsequent analysis. The reconstructed genome of the most recent common ancestor of the MTBC was used as the reference sequence for read mapping (*46*). Trimmed reads were aligned to the ancestral reference genome using BWA (*47*), and SNPs were called using SAMtools (version 1.3.1) (*48*), with a minimum mapping quality of 30. Variants located within repetitive or otherwise poorly mappable genomic regions, including PE/PPE and PE-PGRS family genes, prophage regions, insertion sequences, and other mobile genetic elements, were excluded using a previously described *M. tuberculosis* genome masking scheme (*49*). Premature stop-codon mutations were identified using a custom Perl script by determining whether each nucleotide substitution introduced a stop codon within the annotated coding sequence; substitutions at the native terminal codon and synonymous substitutions within stop codons were not counted. For *pks12*, 7,045 of 12,456 bp (corresponding to 2,336 fully retained codons of 4,152, with a further 25 codons partially masked) were included in the analysis; the remaining 5,411 bp (43.4%) were excluded because they fall within the masked regions, predominantly as regions of reduced pileup mappability with an increased chance for error in interpretation. Functional domains of Pks12 were annotated according to the UniProt entry for the protein. Domain-specific pN/pS ratios were calculated within the same retained intervals using a previously described Python script (*50*).

For each coding sequence, all possible single-nucleotide substitutions were enumerated and classified according to whether they would generate a premature stop codon or a synonymous change, providing gene-specific mutational opportunity counts for the two classes. For each gene under test, genome-wide rates of observed premature stop-codon and synonymous mutations were estimated per mutational opportunity after excluding that gene from the calculation. These leave-one-gene-out background rates were then combined with the gene-specific opportunity counts to calculate the expected proportion, *p*0, of observed mutations in that gene that would be premature stop codons under the null model. Given the total number of observed premature stop-codon and synonymous mutations in the gene, depletion of premature stop codons was tested using a one-sided binomial test, evaluating P(*K* ≤ *k*obs). P values were corrected for multiple testing using the Benjamini– Hochberg procedure.

Following identification of variants for pN/pS and premature stop codon analysis, genes were matched using “Rv” gene IDs with *in vitro* essentiality data from a comprehensive published transposon insertion sequencing (Tn-seq) screen (*19*). *In vitro* essentiality calls were consolidated for simplicity: genes classified as “essential (ES)”, “essential domain (ESD)”, or “growth defect (GD)” in the Tn-seq screen were all considered essential for our analysis, while genes classified as “nonessential (NES)”, “growth advantage (GA)”, or “uncertain” were considered nonessential.

### Bacterial strains and growth conditions

The generation, from *Mycobacterium tuberculosis* (Mtb) H37Rv WT, of the Δ*pks12* and Δ*pks12*::*pks12* (complemented by pYUB2410) strains was described previously (*21*). All knockout and complemented Mtb strains were validated as positive for phthiocerol dimycocerosate (PDIM) by HPLC-ESI-MS of total lipid. All Mtb strains lacking *pks12* carried a hygromycin resistance cassette in its place, and complemented strains carried an integrated kanamycin resistance cassette. Except where otherwise noted, Mtb strains were grown in 7H9 liquid medium with 0.2% glycerol, 10% OADC (BD 212351), and 0.05% tween-80, and stored at -80°C in the same. Hygromycin B (50 μg/ml) and/or kanamycin (20 μg/ml) was added to the medium during maintenance growth, but not during experiments.

For scanning electron microscopy (SEM) experiments, a new set of H37Rv strains was developed in the mc^2^7902 triple-auxotrophic background, which can be used in a biosafety level 2 environment (*51*). *pks12* was deleted from *Mtb* mc^2^7902 using a previously described specialized transduction protocol (*52*). Briefly, the left and right flank of *pks12* were amplified by PCR using primer pairs LL/LR and RL/RR (LL: 5’- TTTTTTTTCCATAAATTGGGATGCGGTCGTGTTGTCAGC-3’, LR: 5’- TTTTTTTTCCATTTCTTGGGGGAGGGCTTCCATTCCTATGATGTCTCACTGA GGTCTCTCTTCGGTCGCATGCTGGAGT-3’; RL: 3’- TTTTTTTTCCATAGATTGGGTTGCTAACATGGTCTCTGCCGAGTGTCTGGTC TCGTAGAGATCGCGGACATGGACCTC-5’, RR: 5’-TTTTTTTTCCATCTTTTGGAATGCGCCGAAACGTCAACT-3’), respectively. The resulting PCR products were digested with *Van91I* and ligated into *Van91I-*digested pYUB1471. The cosmid was digested with *PacI* and ligated into *PacI*-digested shuttle phasmid phAE159. The ligation product was packaged in vitro using GigapackII (Stratagene, La Jolla, CA) and transduced into *E. coli* HB101. The resulting phasmid was electroporated into *Mycobacterium smegmatis* mc^2^155, and a high titer phage lysate was prepared to transduce mc^2^7902. Transductants were screened by three-primer PCR using pks12F (5’- ATGATACCGGGATCGGACAC-3’), pks12R (5’- ATCTCCTCGGCCAACATTCC-3’), and the universal uptag (5’-GATGTCTCACTGAGGTCTCT-3’) incorporated in the LR primer. The primers were designed to generate a 1206-bp amplicon from the wild-type allele and a 751-bp amplicon from the deletion allele. The *pks12* deletion strain was confirmed by whole-genome sequencing. For the mc^2^7902 Δ*pks12*::*pks12* complemented strain, a new complementation vector (pGB55) was generated by cloning *pks12* into the L5-integrating pJEB402 backbone (*53*) under control of the included mycobacterial optimized promoter (MOP). *pks12* and 20 bp of upstream DNA was amplified from Mtb H37Rv genomic DNA in three approximately 4 kb overlapping fragments and assembled into pJEB402 digested with HindIII and EcoRI via Gibson assembly using the HiFi kit (NEB E2621).

*Mycobacterium marinum* M strain and two *pks12* transposon mutant strains, designated *pks12*::Tn- 14377 and *pks12*::Tn-19059, were used for larval and adult zebrafish infections. All carried the plasmid pMSP12:mCerulean (*54*). These strains originated from a sequence-defined *M. marinum* transposon mutant collection provided by C. Cosma and L. Ramakrishnan as described previously (*55*, *56*). The TnMar element present in these mutants encodes hygromycin B resistance. Transposon insertion sites were confirmed by PCR and sequencing, as previously described (*57*). The following primers were used for confirmation: *pks12*::Tn^19059^ F, 5′-AATCGGGTGGTGTCGTGTT-3′, and R, 5′- CAGGAGAACGGCAACACC-3′; and *pks12*::Tn^14377^ F, 5′-GAGTTGTTGGAGCCCTTTGG-3′, and R, 5′-ATCCAACTCTGCTGTGTCCA-3′. *M. marinum* cultures were grown without shaking at 32°C to an optical density (OD600_nm_) of approximately 0.8 in complete 7H9 medium supplemented with 10% OAD (50 g/L bovine serum albumin, 0.5% v/v oleic acid, 20 g/L dextrose, and 8.5 g/L NaCl) and 0.05% Tween 80. When appropriate, hygromycin B (50 μg/ml) or kanamycin (20 μg/ml) was added to the medium. Single-cell aliquots of *M. marinum* fluorescent cultures were prepared for adult and larval zebrafish infections as previously described (*58*) by passage through a 27-gauge needle (BD 309623) followed by filtration through a 5-μm filter (Millex SLSV025LS) and stored at −80°C.

### Mouse infections

6-week old female C57BL/6 mice (Charles River, Kingston, NY)—8 per group, 24 total—were infected via the aerosol route with approximately 100 cfu of the indicated Mtb strains. Mice were maintained with daily health monitoring for 376 days, with euthanasia upon reaching predetermined humane endpoints. Statistics were performed in R using the “survival” package (*59*).

Infections of collaborative cross and other mouse lines were described previously; all data from these infections used in this study were previously published (*6*).

### Zebrafish handling and maintenance

Adult zebrafish were maintained in 3- or 6-liter tanks on a 14-hour light/10-hour dark cycle at 28°C, with water pH maintained between 7.0 and 7.3 and conductivity between 600 and 700 μS. Conductivity was maintained using Instant Ocean Sea Salt (SS15-10), and pH was buffered with sodium bicarbonate (Arm & Hammer Pure Baking Soda, 426292). Adult zebrafish were fed twice daily, once with dry food and once with *Artemia*. Larval zebrafish were maintained at 28.5 °C in 100-mm Petri dishes containing sterile E3 medium (5 mM NaCl, 178 μM KCl, 328 μM CaCl₂, and 400 μM MgCl₂), at a density of 60-100 larvae per dish. For imaging experiments, pigmentation was inhibited by adding PTU (1-phenyl-2-thiourea; Sigma-Aldrich P7629) solution at 1 day post-fertilization.

### Zebrafish infections

Larval zebrafish at 2 days post-fertilization were anesthetized in tricaine at 0.016% final concentration (Syndel MS-222). Approximately 150 fluorescent *M. marinum* bacteria were delivered into the caudal vein of each larva using a borosilicate glass needle. Following injection, larvae were transferred to E3 medium supplemented with PTU for recovery and maintenance. Fluorescent images were captured using a Zeiss Observer Z1 microscope. Fluorescent bacterial burden was quantified at 1 and 5 days post-infection, as previously described (*57*), by quantifying pixels above background using a fixed threshold across samples. Statistical analyses were performed in GraphPad Prism. Representative images were selected based on median fluorescence values.

Adult zebrafish (approximately equal numbers of female and male) were anesthetized with 120 mg/l pharmacological grade tricaine. Single-cell aliquots of *M. marinum* were diluted in 7H9 (Gibco 10010) to generate an inoculum containing approximately 350-400 fluorescent bacteria per 10 µl. Adult zebrafish were injected intraperitoneally with 10 µl of the inoculum using a 27G syringe (BD 08290-3284-38). Following infection, zebrafish were allowed to recover and were maintained in tanks at 28.5°C with daily health monitoring, feeding, and water exchanges. Infected zebrafish were monitored daily and euthanized when they reached predefined humane endpoint criteria approved under the relevant animal care protocol. Statistics were performed in R using the “survival” package (*59*), with zebrafish sex as a covariate.

### Immunogold labeling and transmission EM

For immunogold labeling experiments, virulent Mtb strains grown in tween-containing cultures were washed 3 times with PBS and diluted to an OD_600nm_ of 0.01 in 7H9 media without detergent. They were grown until approximate OD_600nm_ reached ∼1-1.5, as determined by growth of matched tween-containing cultures. Culture media consisted of 7H9 with 0.2% glycerol, with one of three additives at 10% final volume. For standard medium, the additive was OADC (BD 212351). For rich medium, the additive was a form of OADC with additional oleic acid and albumin (“OADC-Hi”): 150 g/L bovine serum albumin, 6.75 mM oleic acid, 20 g/L dextrose, 8.5 g/L NaCl, and 30 mg/L catalase, incubated in glass for at least half an hour at 37°C with shaking, then 0.2 µm filtered to sterilize. For fatty acid-free medium, the additive was “ADN-FAF”: 50 g/L fatty acid-free bovine serum albumin (Thermo J64944.22), 20 g/L dextrose, and 8.5 g/L NaCl, stirred until fully dissolved, then 0.2 µm filtered to sterilize.

After growth, bacteria for TEM were pelleted by 8 minutes of centrifugation at 3500 rpm and resuspended in fixative (PBS with 3% paraformaldehyde and 3% glutaraldehyde, Electron Microscopy Sciences NC1608118) for 24 hours at room temperature. Fixed bacteria were pelleted again, then resuspended in PHEM storage buffer (60 mM PIPES, 25 mM HEPES, 2 mM MgCl2, 10 mM EGTA, pH = 6.9) with 0.5% paraformaldehyde and stored at 4°C.

For immunogold labeling of intact bacteria, fixed Mtb were incubated on 200 mesh formvar-coated hexagonal grids (Electron Microscopy Sciences FF200H-CU-50) for 15 minutes. Thereafter, grids were washed with PBS + 0.02M glycine (Merck K27662101) and incubated with primary antibody CS- 35 at 1:100 dilution (BEI NR-13811) for 45 minutes. Grids were washed 3 times for 5 minutes and incubated with rabbit anti-mouse bridging antibody at 1:200 dilution (DAKO Z0259), then washed another 3 times for 5 minutes. Then, grids were incubated with protein A gold (1:25; Utrecht University), washed 3 times 5 minutes with PBS, and fixed for 5 minutes with 1% glutaraldehyde in PBS. After fixation, grids were washed 3 times for 5 minutes with ddH_2_O and the excess water was removed with blotting paper. Grids were imaged using a Tecnai G2 spirit biotwin transmission electron microscope (ThermoFisher) with Xarosa camera and Emsis software. Images were analyzed using ImageJ with FIJI plugins.

For cryosectioning and immunogold labeling of sections, fixed Mtb were washed 3 times with PBS + 0.02M glycine, after every wash the Mtb were centrifuged at 2730 *x g* and the supernatant was removed. Thereafter, the Mtb were incubated with 12% gelatin (Sigma G2500-500G) for 5 minutes at 37°C and centrifuged at 10950 *x g*. Samples were incubated on ice to solidify the gelatin and 1x1mm blocks were cut using a razor blade. These gelatin blocks were incubated overnight in 2.3M sucrose at 4°C while rotating. The next day, each gelatin block were placed on a metal pin (Leica, 16701950) and snap frozen using liquid nitrogen. Samples were stored in liquid nitrogen until sectioning. Samples were sectioned at -120°C using a cryo-ultra microtome (Leica Ultracut UC6), with 60 nm sections cut using a diamond knife (Diatome). Sections were transferred to 200 mesh hexagonal copper grid (EMS, FF200H-CU-50) using a 1:1 mixture of methylcellulose (Sigma M6385-250G) and 2.3M sucrose (Merck 7653.5000). Grids were stored at 4°C.

Cryosectioned grids were incubated on a 2% gelatin plate for 30 minutes at 37°C. Thereafter, grids were washed 5 times for 2 minutes with PBS + 0.02M glycine, blocked with 1% BSA in PBS + 0.02M glycine for 3 minutes, and incubated with primary antibody CS-35 (1:100; BEI resources NR-13811) in 1% BSA in PBS for 45 minutes. Grids were washed 5 times for 2 minutes with PBS + 0.02M glycine, blocked with 0.1% BSA in PBS + 0.02M glycine, and incubated with rabbit anti-mouse bridging antibody at 1:200 dilution (DAKO Z0259) for 20 minutes. Grids were washed 6 times for 3 minutes with PBS + 0.02M glycine, blocked with 0.1% BSA in PBS + 0.02M glycine, and incubated with protein A gold (1:25, Utrecht University) for 20 minutes. Grids were washed 6 times for 3 minutes with PBS and fixed with 1% glutaraldehyde for 5 minutes. Grids were washed 10 times for 2 minutes with ddH_2_O and stained with uranylacetate in methylcellulose for 5 minutes in the dark. Grids were blotted and dried at room temperature. Grids were stored at room temperature until imaging. Samples were imaged using a Tecnai G2 spirit biotwin transmission electron microscope (ThermoFisher) with a Xarosa camera and Emsis software. Images were analyzed using ImageJ with FIJI plugins.

### Transmission and scanning EM without immunogold

Mtb mc^2^7902 strains were cultured in 7H9 medium (Sigma M0178-500g) supplemented with 1% ADC (BD 211887), PMLA additive (final concentration of 24 µg/mL pantothenate, 50 µg/mL methionine, 60 µg/mL leucine, 200 µg/mL arginine), and with or without 0.05% Tween-80 (Sigma P8074). Cultures were incubated at 37°C and shaken at 50 rpm. After 8 days, the cultures were fixed in PHEM buffer with 2% paraformaldehyde and 0.2% glutaraldehyde. After 24 hours, fixative was replaced with storage buffer (0.64% paraformaldehyde in 0.1M phosphate buffer) and bacteria were embedded for transmission electron microscopy immediately.

For transmission EM to observe capsular organization, fixed mc^2^7902 cultures were washed with ddH_2_O and incubated with 1% osmium tetroxide with or without 1.5% potassium ferrocyanide (Merck 4973) for 1 hour. Samples were washed with ddH_2_O and dehydrated in an increasing percentage of ethanol (70-100%). Thereafter, samples were incubated with propylene oxide (Sigma 8.07027.1000) for 45 minutes, 1:1 propylene oxide and Epon resin for 2 hours, and 1:2 propylene oxide and Epon resin overnight. The next day, samples were impregnated with Epon resin at 37°C for 4 hours and Epon was refreshed after 4 hours and incubated for a further 3 hours at 37°C. Thereafter, samples were polymerized at 65°C for 3 days and sectioned on a ultramicrotome (Leica Ultracut UC6 ultramicrotome) in 60 nm sections using a diamond knife (Drukker), then were mounted on a formvar-coated 200 mesh hexagonal grid (Electron Microscopy Sciences FF200H-CU-50). Grids were stained using 3.5% uranyl acetate for 5 minutes, washed with ddH_2_O, incubated with lead citrate (EMS, 22410) for 3 minutes, then washed and blotted with blotting paper. Samples were imaged on a Tecnai G2 spirit biotwin transmission electron microscope (ThermoFisher) with a Xarosa camera and Emsis software.

For scanning EM, fixed Mtb mc^2^7902 cultures were washed with ddH_2_O for 10 minutes and incubated with 1% osmium tetroxide and 1.5% potassium ferrocyanide for 1 hour. Samples were washed with ddH_2_O and dehydrated in an increasing concentration of ethanol (30-100%). Samples are incubated on a poly-L-lysine coated glass slide and placed in a critical point dryer (Leice CPD 300) with the following settings: (CO2 in: slow, exchange speed 3 with 14 cycles and gas out heat slow/ speed slow). Samples were sputter coated (Leica ACE600) with 6 nanometer platinum/palladium. Samples were imaged using a Zeiss Gemini scanning microscope at 3kV with an Inlense detector.

### EM statistics

Graphpad prism versions 10.6.0 and 11.0.2 were used for specific statistical analyses. First a Shapiro-Wilk test (10.6.0) was performed to determine whether the data were normally distributed. Data for electron-dense structures and CS-35 immunogold differential localization were normally distributed; ordinary one-way ANOVAs with Dunnett’s multiple comparisons test were performed (11.0.2). Data for the presence of indentations were not normally distributed, a Kruskal-Wallis test with Dunn’s multiple comparisons test was performed (11.0.2). For CS-35 immunogold abundance on the surface and inner membrane, statistics were performed using with linear mixed-effects models in R, using the “lme4”, “lmertest”, and “emmeans” packages.

### Lipid preparation

Before culture for lipid extraction, bacterial pellets were washed three times with PBS, then grown in media lacking detergent. Mtb was grown in 7H9 with 0.2% glycerol and 10% OADC (BD 212351). *M. marinum* was grown in 7H9 with 0.2% glycerol and 10% ADN (50 g/L bovine serum albumin, 20 g/L dextrose, 8.5 g/L NaCl). Cultures were started at an OD_600nm_ of 0.005 at 15-40 mL scale, then grown until matched tween-containing cultures reached an OD_600nm_ of 0.6-0.8 (for Mtb) or 1.0-1.5 (for *M. marinum*). Bacteria were pelleted for 10 minutes at 3500 rpm, then washed 2 times with Optima grade water. After the second wash, bacteria were pelleted again and resuspended in 1 mL methanol, then transferred without pipet tip contact to a chloroform:methanol mixture to form a final ratio of 1:2 chloroform:methanol. This mixture was inverted to mix and left overnight at room temperature to sterilize. All steps following first chloroform contact were performed exclusively with glass tubes and pipets, using Optima grade solvents.

After sterilization and at least one hour of mixing on an orbital shaker, samples in chloroform:methanol were pelleted by centrifugation, and supernatants were collected. Pellets were incubated further in 1:1 chloroform:methanol, then in 2:1 chloroform:methanol, each for at least an hour on an orbital shaker. After each step, samples were pelleted and supernatants were collected. Supernatants for each sample were combined (∼12 mL total per sample) and evaporated using a GeneVac. Evaporated supernatants were then re-solvated in 3 mL 1:1 chloroform:methanol, sonicated using standard settings (Branson 5510 water bath), and pelleted for 30 minutes at 3500 rpm. Fully clarified supernatants were transferred to pre-weighed 4 mL glass vials and evaporated using a GeneVac. Evaporated lipids were weighed, re-dissolved in 1:1 chloroform:methanol to 1-2 mg/mL, and stored sealed in the dark at -20°C.

Three-phase separation of total Mtb lipids was performed based on a published method (*60*). Total Mtb lipids (100 µg) were evaporated into glass centrifuge tubes under nitrogen. An HPLC-clean four- solvent mix was added to the tubes: 1 mL hexanes, 1 mL methyl acetate, 0.75 mL acetonitrile, and 1 mL water. Samples were vortexed for ∼5 seconds, sonicated, and incubated for at least an hour on an orbital shaker. Tubes were centrifuged at 3500 rpm for 10 minutes to form clean separation of 3 phases. Each phase was removed in turn from the bottom of the tube using a Hamilton glass syringe with a long stainless steel needle. The syringe was washed thoroughly in fresh isopropanol in between phases. Separated phases in glass vials were evaporated using a GeneVac, and either analyzed directly or saponified as described below.

For HPLC-ESI-MS analysis of total lipids, samples (50-100 µg) were evaporated under nitrogen in glass vials using an N-evap and re-solvated in starting solvent for HPLC. For analysis of saponified lipids, samples were evaporated under nitrogen in glass vials, then re-solvated in 50 µL methanol. Saponification was carried out by adding 5.55 µL 1N NaOH and incubating samples for at least 2 hours at 37°C. After saponification, NaOH was neutralized by adding 5.55 µL 1M formic acid in water. Saponified samples were evaporated under nitrogen, redissolved in 50 µL 1:1 chloroform:methanol and transferred to remove excess salts, then evaporated again under nitrogen and re-solvated in starting solvent for HPLC.

### Lipid analysis

Total lipids and saponified lipids were analyzed by reversed phase HPLC-ESI-MS, as described (*17*). Briefly, following sample solvation in solvent A (95:5 methanol:water with 2 mM ammonium formate), 10 μL (10 μg) lipid was injected per sample. Analysis was performed using an Agilent Poroshell 120 EC-C18 column (1.9 μm particle size, 3.0 x 50 mm, 699675-302) on an Agilent 1260 infinity HPLC, with an Agilent 6230 electrospray ionization time-of-flight (ESI-TOF) mass spectrometer. HPLC flow rate was 0.15 mL/minute, with the following ratios of solvent A to solvent B (90:10:0.1 1- propanol:cyclohexane:water with 3 mM ammonium formate): 0-4 minutes (100% A); 4-10 minutes (gradient to 100% B); 10-15 minutes (100% B); 15-20 minutes (gradient to 100% A); 5-minute postrun (100% A). Additional settings: 325°C gas temperature, 8 L/minute drying gas flow rate, 35 PSI nebulizer pressure, and 3500 V. Spectra were collected at 2 spectra/second from 100 to 3200 *m/z*, in the negative ion mode.

Total lipids were analyzed by normal phase HPLC-ESI-MS, as described (*32*). Briefly, following sample solvation in solvent A (70:30 hexanes:isopropanol with 0.1% formic acid and 0.05% NH_4_OH), 20 μL (20 μg) lipid was injected per sample. Analysis was performed using an Inertsil Diol 120 silica column (3 μm particle size, 2.1 x 150 mm, 5020-05415) on an Agilent 1260 infinity HPLC, with an Agilent 6530 electrospray ionization quantitative time-of-flight (ESI-QTOF) mass spectrometer or an Agilent 6230 electrospray ionization time-of-flight (ESI-TOF) mass spectrometer. HPLC flow rate was 0.15 mL/minute, with the following ratios of solvent A to solvent B (30:70 methanol:isopropanol with 0.1% formic acid and 0.05% NH_4_OH): 0-10 minutes (100% A); 10-17 minutes (gradient to 50% B); 17-22 minutes (50% B); 22-30 minutes (gradient to 100% B); 30-40 minutes (gradient to 100% A); 40-44 minutes (100% A); 5-minute postrun (100% A). Additional settings: 325°C gas temperature, 5 L/minute drying gas flow rate, 15 PSI nebulizer pressure, and 3500 V. Spectra were collected at 1.02 spectra/second from 100 to 3200 *m/z*, in the negative and positive ion modes in separate runs.

For fractionation of total lipids into 2-minute bins prior to fraction saponification, normal phase HPLC was carried out as described, with the LC flow redirected to glass vials rather than the mass spectrometer, then evaporated under nitrogen. For CID-MS of MPMMM, normal phase HPLC-ESI-MS was carried out as described, with 100V of collision energy directed at a medium (4 *m/z*) mass window starting at *m/z* 1826, for the first 10 minutes of the run. MS/MS data were collected from 40-3000 *m/z*.

Untargeted lipidomic analysis was performed on normal phase data in the negative mode. Agilent .d files were converted to .mzXML files using msConvert (ProteoWizard) and uploaded for peak picking to XCMSOnline. Peak picking was performed using the following settings: centWave feature detection (10 ppm, snthr=5, peakwidth=20-120, mzdiff=-0.001), obiwarp retention time correction (profStep=0.5), and density grouping (bw=5, mzwid=0.01, minfrac=0.3). Statistical analysis was performed using LIMMS in R, with default LIMMA settings for imputation of zeroes, normalization, and testing (*36*). Ion peaks were matched within 10 ppm and censored and annotated using the MycoLOBSTAHs database of theoretical mycobacterial lipids, and further annotated using the MycoMass database for known Mtb lipids (*32*, *37*).

Targeted lipid analysis was performed using Agilent Masshunter Qualitative Analysis software. Ion chromatograms were extracted from centroid data with a 10 ppm symmetric mass window, and smoothed with default settings. Mass spectra were extracted from profile data.

### Mycolyltransferase assays

Synthesis of pure MPM substrate was previously described (*61*). TMM substrate was purified from total mycobacterial lipids, prepared as described above in “Lipid preparation”, from Mtb Δ*pks12* and *Mycobacterium smegmatis* grown in standard media. Total mycobacterial lipid (>5 mg) was spotted in a line onto a 500 µm thick 20 cm x 20 cm silica gel plate, pre-cleared with 60:30:6 chloroform:methanol:water. Lipids were separated with a 60:16:2 chloroform:methanol:water mobile phase, then the plate was thoroughly dried. The retention factor (Rf) for TMM was determined based on a pre-purified TMM standard, run alongside the preparative sample, then cleaved from the rest of the plate and visualized by spraying to saturation with acidic cupric acetate (8% v/v H_3_PO_4_, 3% w/v Cu[CH_3_COO]_2_) and oven-heating until lipid spots were visually charred (approximately 30 minutes). Silica gel at the TMM Rf was scraped and extracted with 1:1 chloroform:methanol, followed by pelleting and isolation of the supernatant to yield pure TMM. TMM was evaporated in a pre-weighed glass vial and weighed, then re-solvated in 1:1 chloroform:methanol to 1 mg/mL. TMM identity and purity was confirmed via HPLC-ESI-MS analysis.

Antigen 85A and Antigen 85C were expressed and purified according to a published protocol (*62*), with minor deviations. Briefly, pET23b-fpbA and pET23b-fbpC2 were transformed into E. coli C41 (DE3) cells. Cells were cultured in terrific broth with 50 μg/mL Kanamycin. Recombinant protein expression was induced with 1 mM IPTG, and the cells were cultured overnight at 16°C before harvesting the following day. Cell pellets were resuspended in purification buffer (50 mM sodium phosphate, pH 7.5, 500 mM NaCl) containing 25 mM Imidazole, and then lysed by sonication. Cell lysate was clarified through centrifugation at 18,000 rpm. The recombinant protein was purified by Imobilised Metal Affinity Chromatography (IMAC); clarified lysate was applied to a nickel column and the recombinant protein was washed and eluted in purification buffer with increasing concentrations of imidazole. The fractions containing purified recombinant protein (as analysed by SDS-PAGE) were dialysed against 50 mM Tris, pH 7.5, 500 mM NaCl, 10 % glycerol, before concentrating and freezing at -20°C in aliquots.

TMM (50 µg) and MPM (50 µg) were combined in glass vials and evaporated under nitrogen, with TMM-only vials as controls. 100 μL of 50 mM sodium phosphate (pH 7.5) with 20-40 μM FbpA or FpbC was added, with no enzyme vials as controls. Vials were incubated at 37°C for 1 hour. Addition of 6 mL chloroform:methanol:water (10:10:3) quenched the reaction. A further 1.125 mL water and 2.625 mL chloroform was added to the glass vials, which were then vortexed and centrifuged to separate the organic and inorganic layers. The organic lower layer was taken into a new glass tube and 3 mL chloroform:methanol:water (3:47:48) was added, vortexed, centrifuged and the upper top layer was discarded. This step was repeated. The lower layer was transferred into a clean tube and dried for mass spectrometry analysis.

Mycolyltransferase assay products were analyzed by normal phase HPLC-ESI-MS, as described above.

### MPMMM model diagram creation

For the model diagram of MPMMM biosynthesis and presumed localization, membranes and capsular contents were adapted with permission from published figures (*23*, *37*).

---

**Fig. S1.**
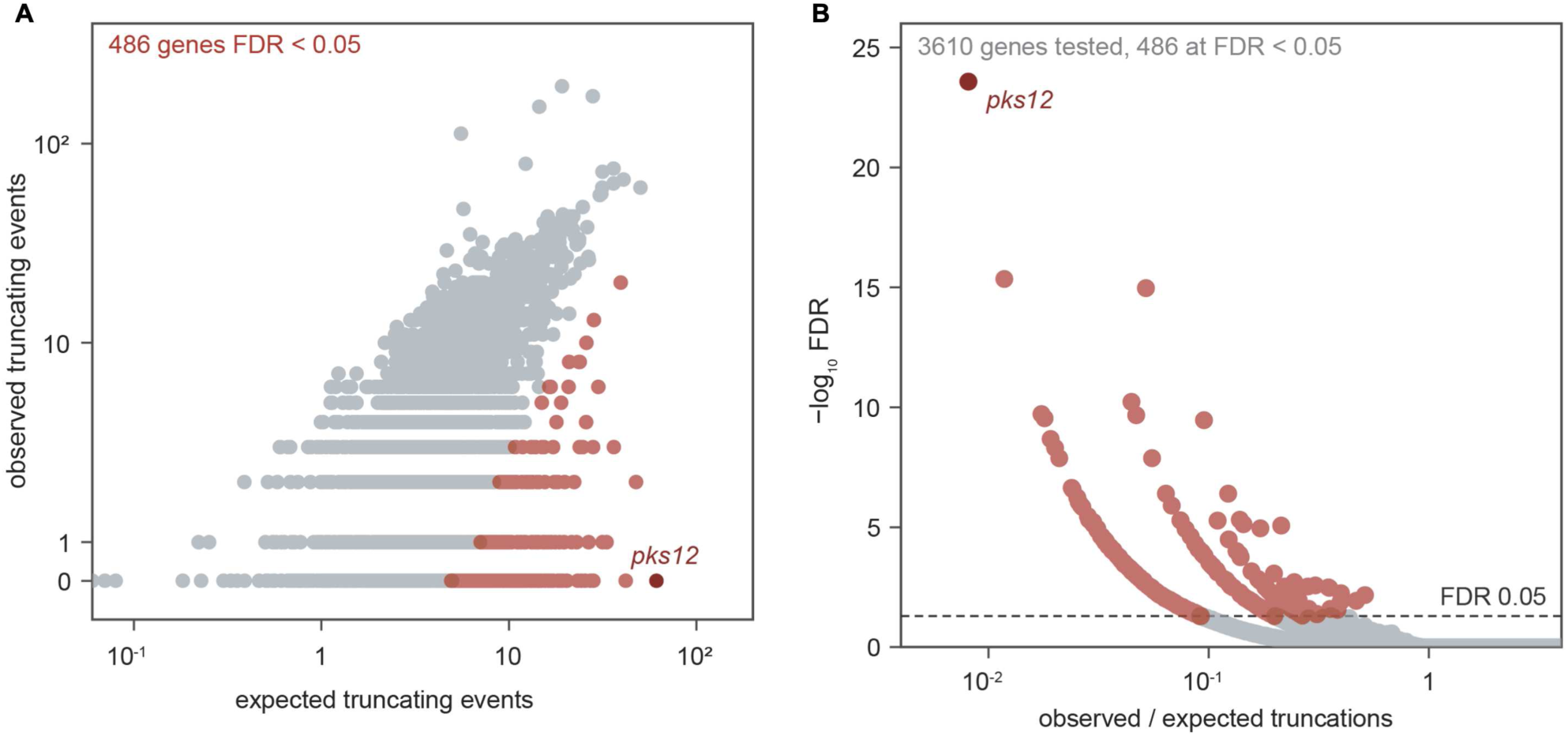
***pks12* is the most significantly under-truncated gene in Mtb.** (**A**) Anticipated versus measured gene truncations via premature stop codon mutation, for all mappable genes in the Mtb genome, with expectations set by gene-specific mutational opportunity and measured genome-wide mutation rates. (**B**) Significance of lower-than-expected truncation rates as measured by false discovery rate (FDR) following Benjamini-Hochberg correction.

**Fig. S2.**
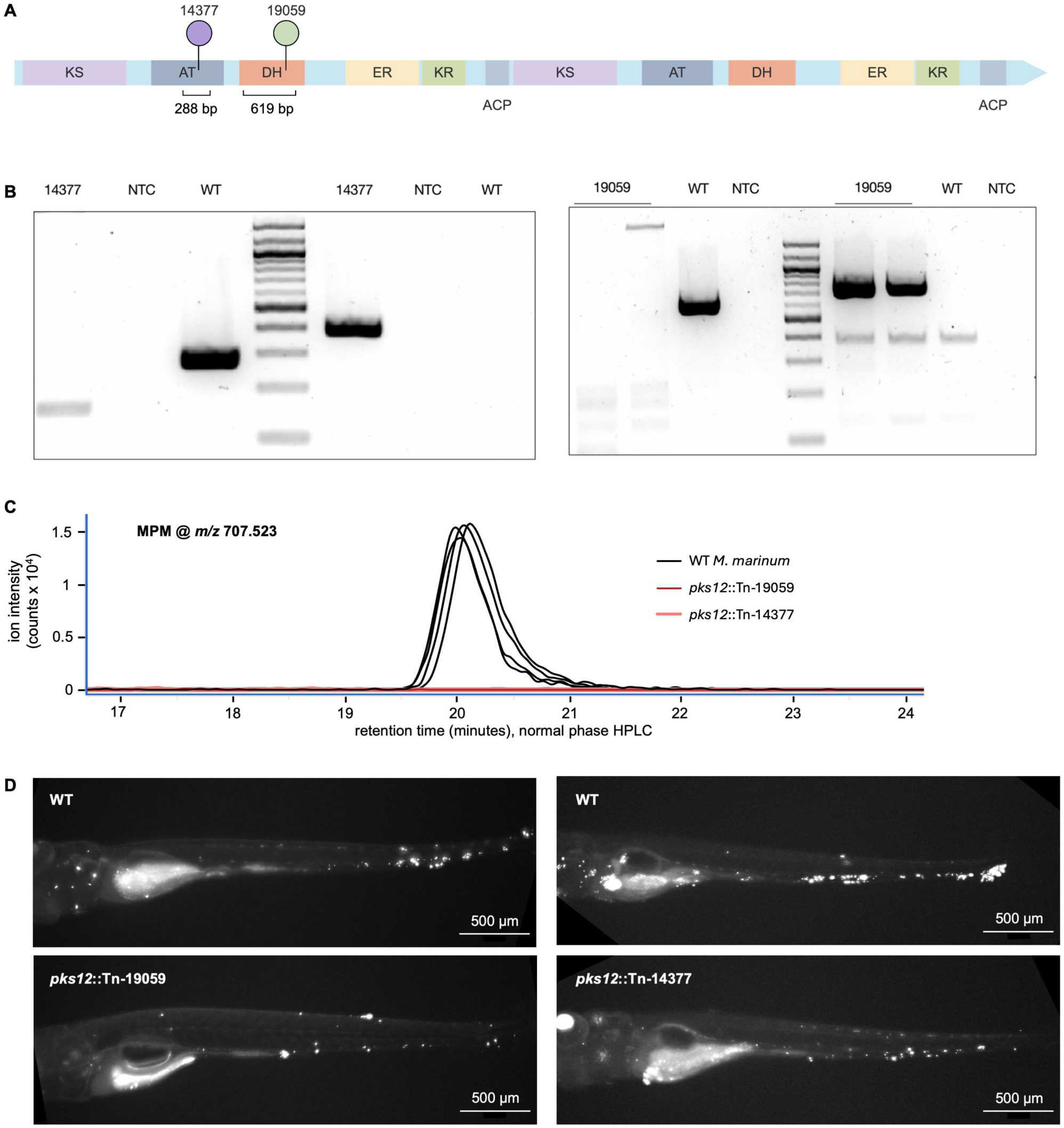
**Analysis of *pks12* transposon mutants in *M. marinum***. (**A**) Schematic representation of the *M. marinum* MMAR_3025/*pks12* coding sequence, showing predicted protein domains and the positions of two independent transposon insertions, *pks12*::Tn-14377 and *pks12*::Tn-19059. Domains are labeled as ketoacyl synthase (KS), acyltransferase (AT), dehydratase (DH), enoyl reductase (ER), ketoreductase (KR), and acyl carrier protein (ACP). The schematic was generated using the IBS 2.0 web server ^(ref)^. (**B**) PCR analysis of *pks12*::Tn-14377 and *pks12*::Tn-19059 compared with WT, no-template control (NTC), and TriDye 100 bp DNA Ladder (New England Biolabs N3271S). WT PCR amplicon sizes ran as expected compared to a DNA ladder: 288 bp for the *pks12*::Tn-14377 locus and 619 bp for the *pks12*::Tn-19059 locus. Mutant amplicons are expected to be approximately 100 bp larger because of the transposon insertion. (**C**) High performance liquid chromatography-mass spectrometry (HPLC-MS) chromatogram for the Pks12 lipid product mannosyl phosphomycoketide (MPM) in WT and *pks12*::Tn strains. (**D**) Representative fluorescence images at 5 days post-infection of zebrafish larvae infected with an initial dose of 150–250 fluorescent bacilli, shown for *pks12*::Tn-14377 and *pks12*::Tn-19059 mutants and intra-experimental matched WT animals.

**Fig. S3.**
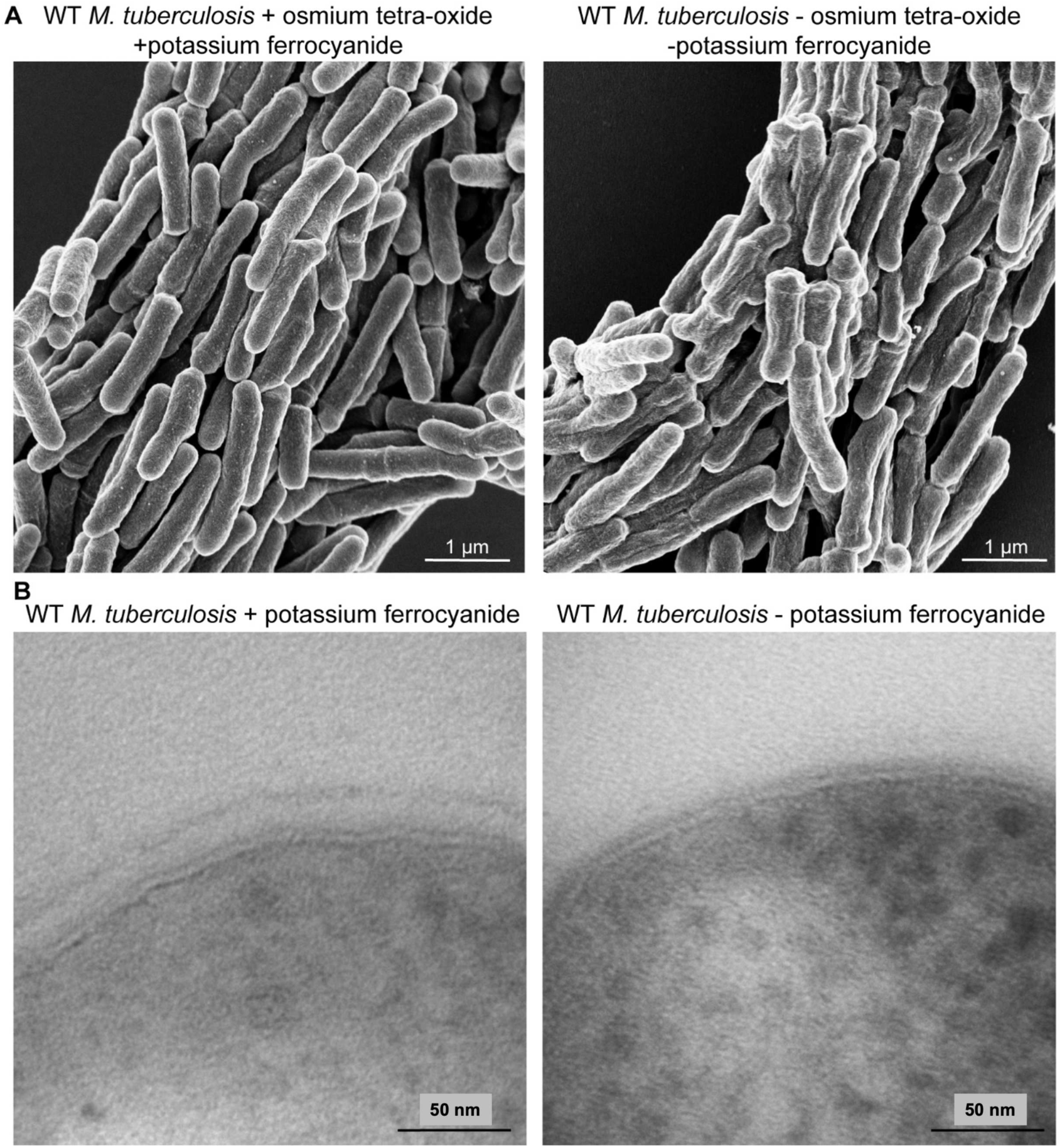
Potassium ferrocyanide improves Mtb structure in SEM and shows the capsular layer in TEM. (**A**) Representative SEM images of WT Mtb with and without osmium tetra-oxide (osmium tetroxide), showing preservation of intact surface morphology. (**B**) Representative TEM images of WT Mtb embedded with or without potassium ferrocyanide, showing preservation of the capsular glycan layer.

**Fig. S4.**
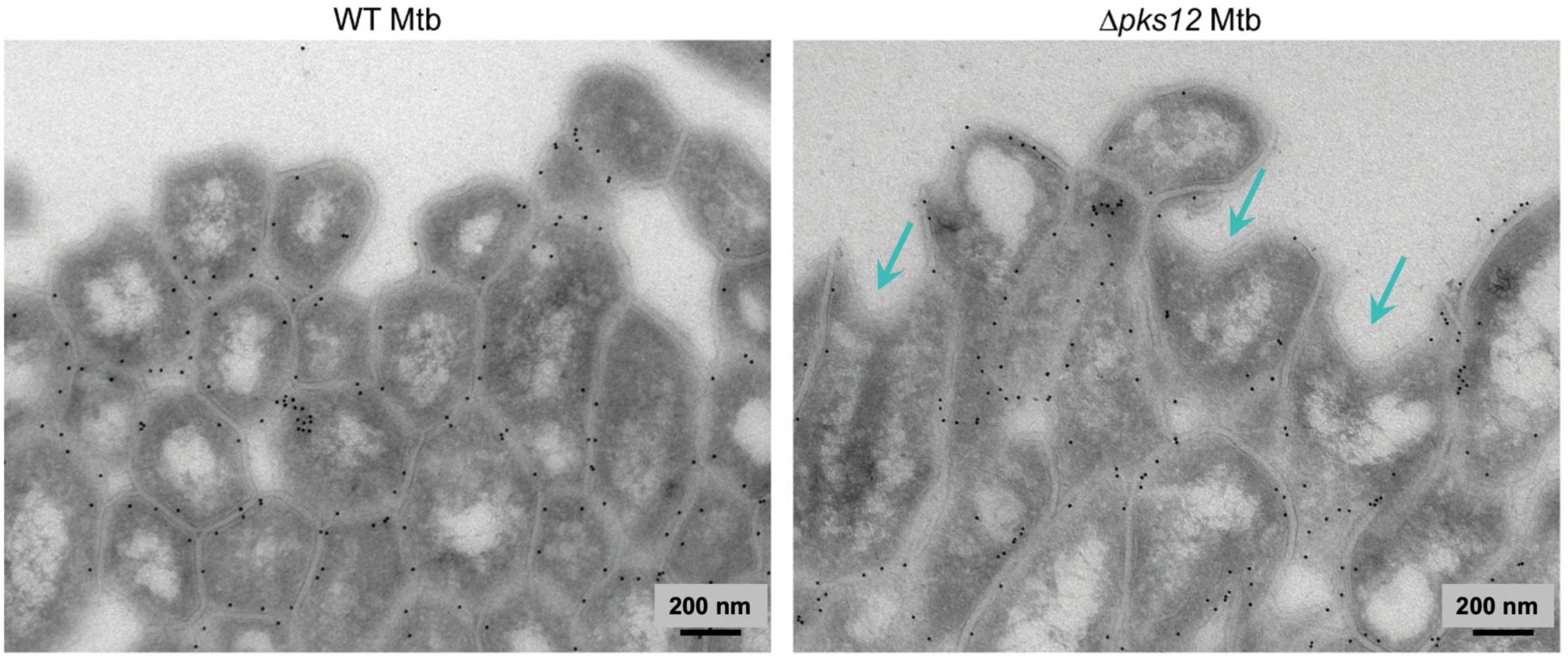
Indentations in Δ*pks12* Mtb extend to all layers of the cell envelope. Representative TEM images of cryo-sections of *M. tuberculosis* cords of WT Mtb (left) and Δ*pks12* Mtb (right). Cyan arrows indicate bacterial indentations. Sections were labelled with CS-35 antibody immunogold.

**Fig. S5.**
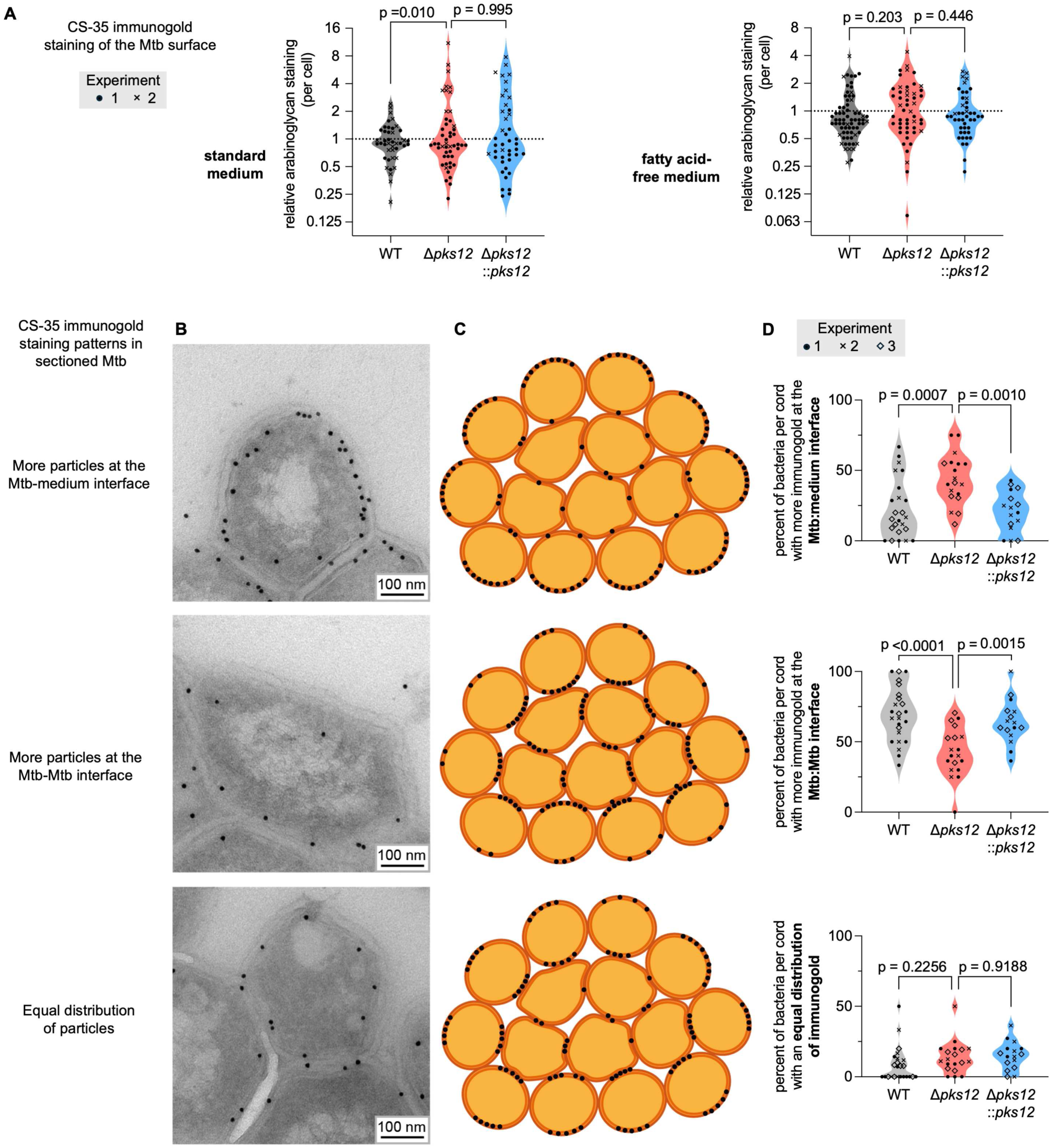
Effects of deleting *pks12* on arabinoglycan surface exposure and distribution. (**A**) Quantification of of CS-35 immunogold staining for surface arabinoglycans on intact WT, Δ*pks12,* and complemented Mtb, grown in different media. Data points represent individual bacteria across two independent experiments, indicated by different symbols; statistics were performed using linear mixed-effects models with Tukey’s multiple comparisons test. (**B**) Representative TEM images of bacteria with differential localization of CS-35 immunogold between the Mtb-Mtb cellular interface and the Mtb-medium interface. (**C**) Schematic representation of Mtb within a cord (cross-sectioned) with differential localization of CS-35 immunogold, as described in (B); orange represents the cytosol and dark orange the cell envelope, while black dots represent CS-35 immunogold. (**D**) Quantification of the percentage of Mtb per cord with with differential localization of CS-35 immunogold. Data points represent cords from 3 individual experiments, indicated by different symbols; statistics were performed using an ordinary one-way ANOVA with Dunnett’s multiple comparisons test.

**Fig. S6.**
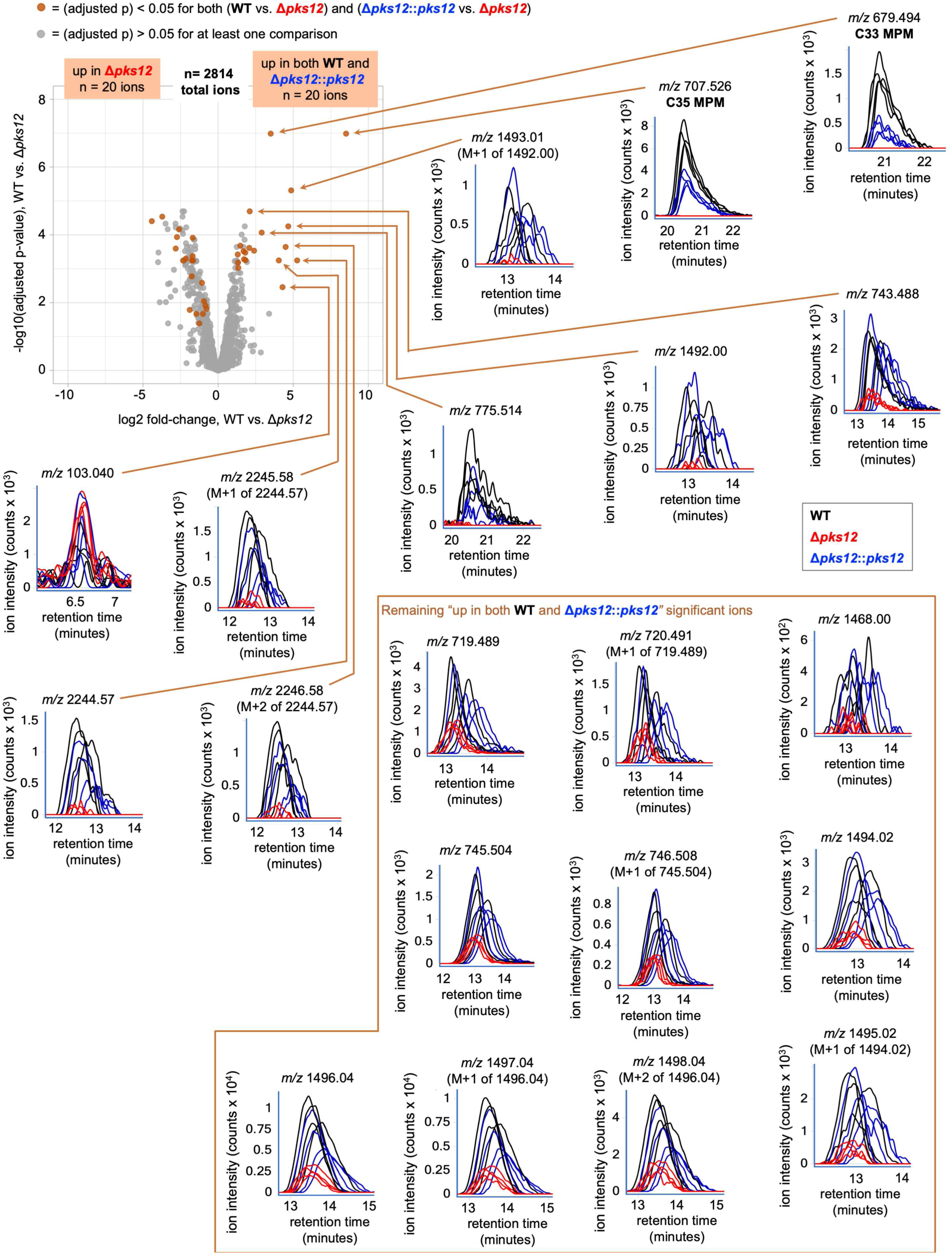
Lipidomics with automated peak picking does not identify any non-MPM *pks12* products. Lipidomic comparison of WT and complemented (Δ*pks12*::*pks12*) Mtb with Δ*pks12* Mtb; volcano plot for WT vs. Δ*pks12* shown (top left). Chromatograms are shown for all 20 ions that are significantly lower (adjusted p < 0.05) in the Δ*pks12* mutant for both comparisons (WT vs. Δ*pks12* and Δ*pks12*::*pks12* vs. Δ*pks12*). Ions are labeled with XCMS-calculated m/z values; it is noted when they are derived from known lipids or when they are ^13^C isotopes (M+n) of other significant ions. Only the two MPM ions with completely flat Δ*pks12* (red) chromatograms are unambiguously and exclusively produced downstream of *pks12*.

**Fig. S7.**
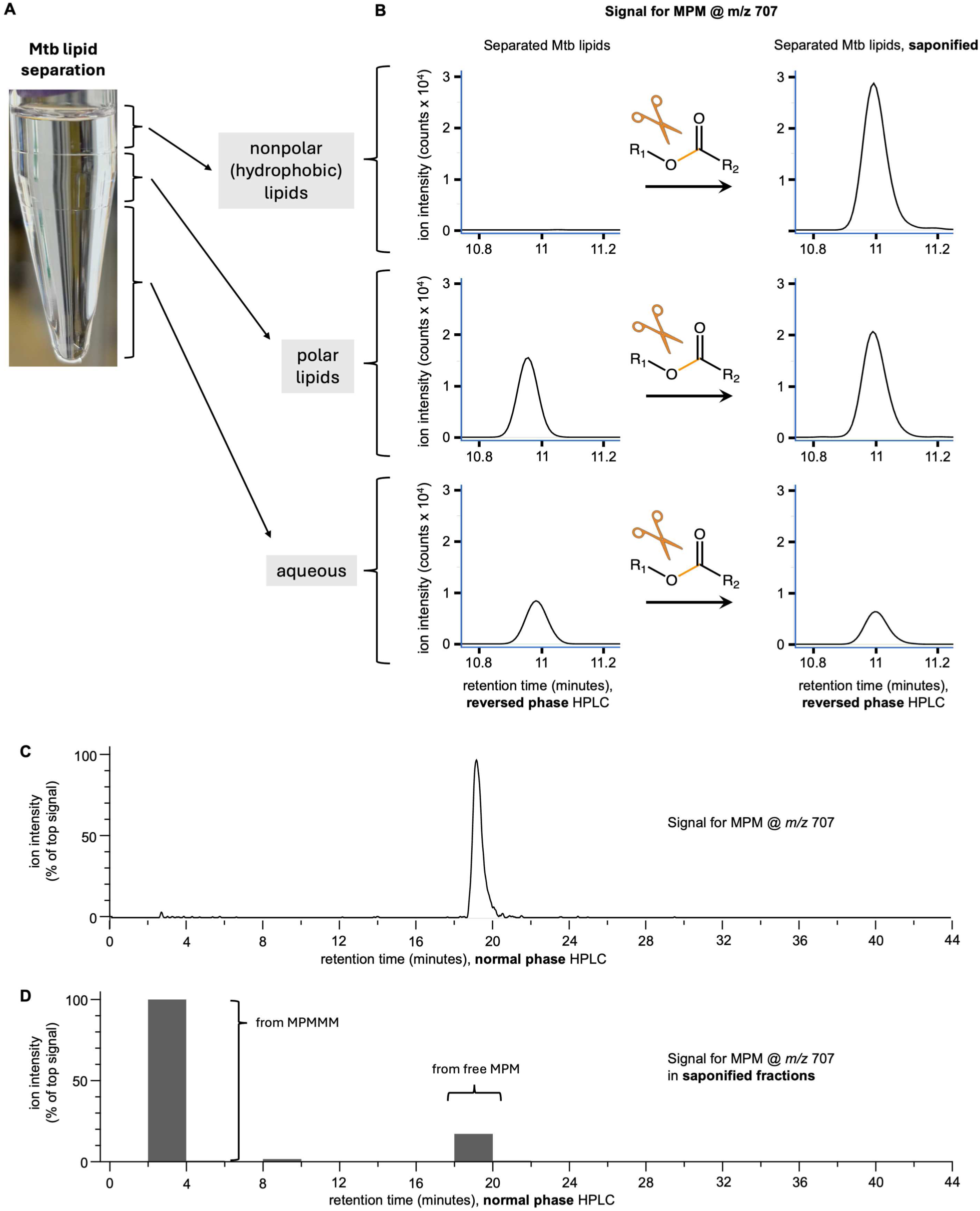
Saponifiable MPM is released from hydrophobic material. (**A**) 3-phase method used to separate Mtb lipids. (**B**) Targeted analysis of the 3 separated phases of Mtb lipids for MPM (by reversed phase HPLC-ESI-MS) before (left) and after saponification (right). (**C**) Normal phase HPLC-ESI-MS analysis of Mtb total lipids, with targeted analysis for MPM. (**D**) Fractionation of Mtb total lipids by normal phase HPLC with collection every two minutes, followed by saponification, and then targeted analysis for MPM (by reversed phase HPLC-ESI-MS). Data shown are representative of three independent experiments.

**Fig. S8.**
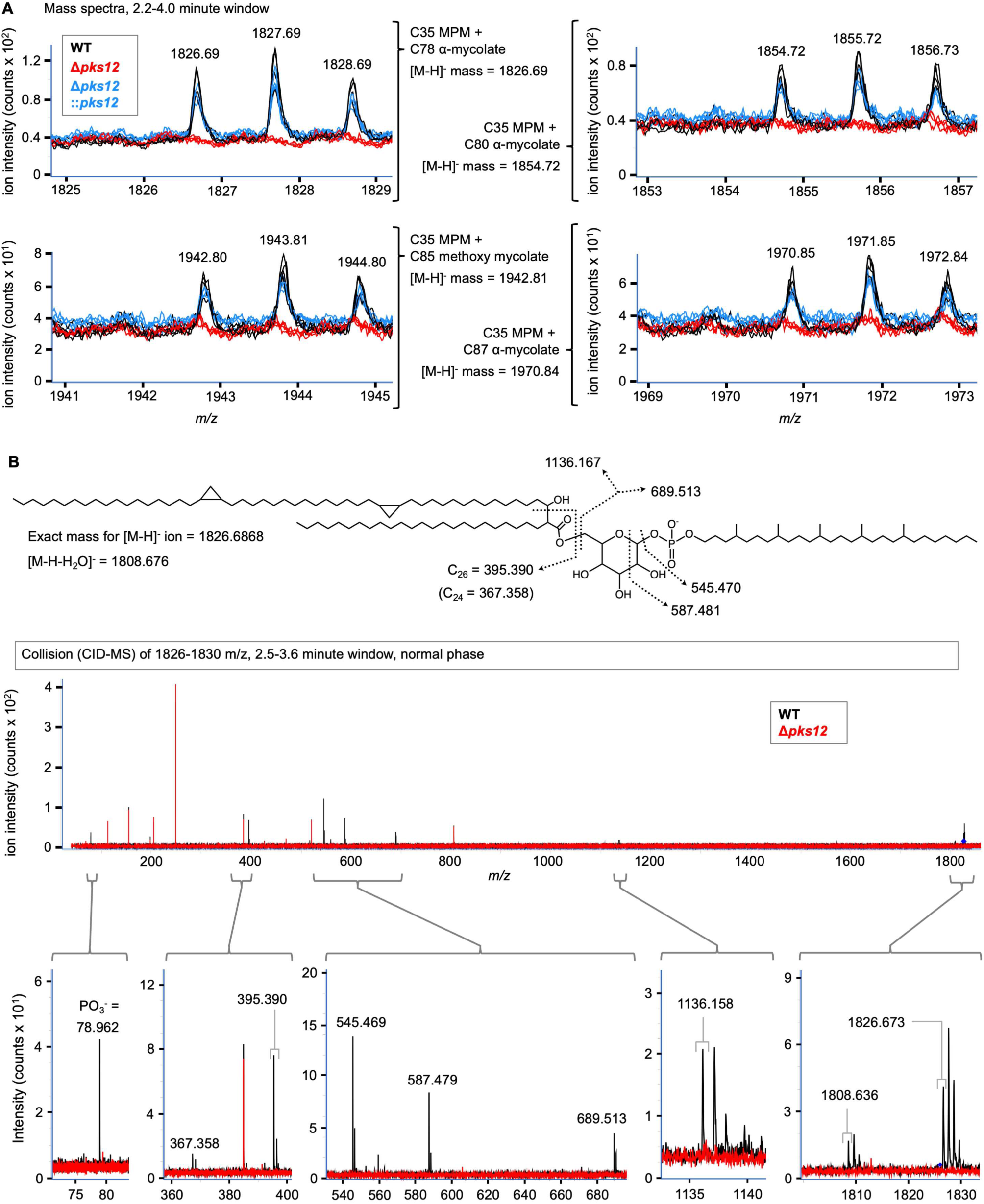
A mycolated form of MPM, MPMMM, is *pks12*-dependent. (**A**) HPLC-ESI-MS spectra for four MPMMM molecules with differing mycolyl chain length. (**B**) Collisional mass spectrometry (CID-MS) for the most abundant form of MPM at *m/z* 1826 (including abundant ^13^C isotopes), with Δ*pks12* CID-MS products used to filter out nonspecific (red) fragment ion peaks. All peak *m/z* values were assigned by MassHunter software, all ion mass values were calculated from chemical formulae.

**Fig. S9.**
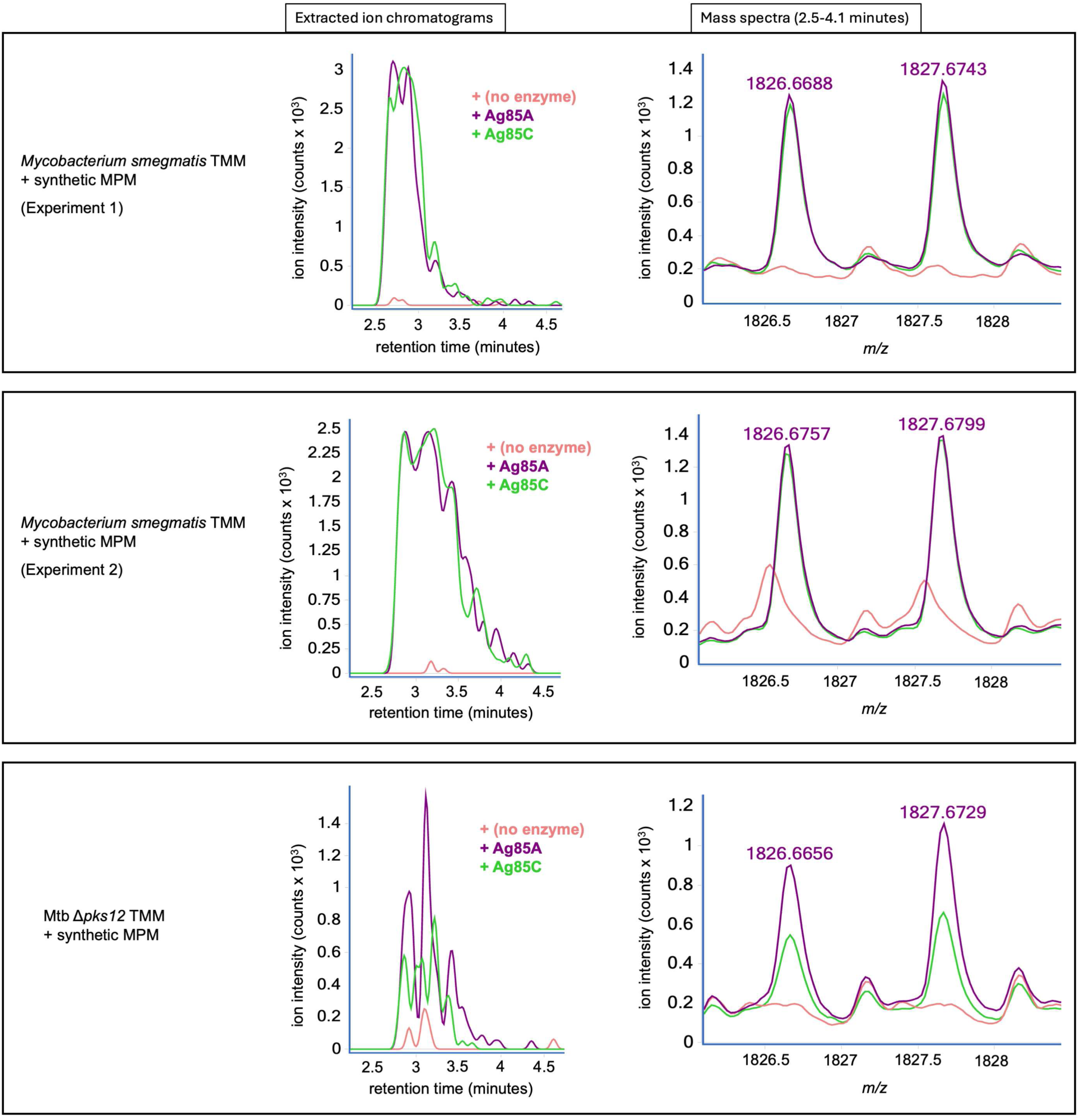
Ag85 enzymes synthesize MPMMM across TMM preparations and independent experiments. HPLC-ESI-MS chromatograms (left) of the monoisotopic ion of MPMMM @ 1826 *m/z*, for products of *in vitro* reactions with or without Ag85 enzyme. Mass spectra (right) show each monoisotopic [M-H]- and the second isotope (single-^13^C-containing) peak.

